# Autism risk genes converge on *PBX1* to govern neural cell growth

**DOI:** 10.1101/2025.03.12.642693

**Authors:** Shuai Fu, Anthony Wynshaw-Boris

## Abstract

The alteration of neural progenitor cell (NPC) proliferation underlies autism spectrum disorders (ASD). It remains unclear whether targeting convergent downstream targets among mutations from different genes and individuals can rescue this alteration. We identified *PBX1* as a convergent target of three autism risk genes: *CTNNB1*, *PTEN*, and *DVL3*, using isogenic iPSC-derived 2D NPCs. Overexpression of the PBX1a isoform effectively rescued increased NPC proliferation in all three isogenic ASD-related variants. Dysregulation of *PBX1* in NPCs was further confirmed in publicly available datasets from other models of ASD. These findings spotlight *PBX1*, known to play important roles during olfactory bulb/adult neurogenesis and in multiple cancers, as an unexpected and key downstream target, influencing NPC proliferation in ASD and neurodevelopmental syndromes.

## Main Text

ASDs are a group of highly heterogenous neurodevelopmental disorders characterized by core symptoms, including impaired social interaction and repetitive behaviors^1^. Alterations in cortical neurogenesis and neuronal function are implicated in ASD pathology^2–5^. Increased NPC proliferation has been found in individuals with ASD, particularly those exhibiting early brain overgrowth^6,7^. Variants in ASD risk genes have been linked to this hyperproliferation phenotype^4^. However, whether mutations in different high-confidence ASD risk genes converge on a shared downstream target to drive increased NPC proliferation remains unexplored.

In this study, we used 2D NPC models to investigate the effects of patient-specific mutations in three ASD risk genes (*CTNNB1*, *PTEN*, and *DVL3*) each modeled individually within the same control iPSC line, focusing on phenotypic changes, gene expression, and alternative splicing. Mutations of these genes are associated with ASD and early brain growth^4,6,8–12^. We found that mutations in these genes lead to increased NPC proliferation and converge on dysregulation of PBX1, a downstream transcription factor critical for cellular proliferation and differentiation.

PBX1, known for its role in regulating proliferation across several cancer types^13–16^, exists in two main isoforms, PBX1a and PBX1b. Inclusion of exon 7, which is conserved across mammals^17^, resulted in PBX1a. During development, Pbx1b is mostly expressed in early embryonic tissues, whereas Pbx1a is found in neural tissues, expressed in both progenitors and neurons, contributing to cortical patterning and dorsal-ventral organization during embryogenesis^18^. During adult neurogenesis, PBX1 is expressed in subventricular zone (SVZ) progenitors^19^ and olfactory bulb^20^. PBX1 alternative splicing is also important in neuronal differentiation. The splicing factor Ptbp1 suppresses Pbx1a isoform expression by skipping exon 7 in the mESC, removal of Ptbp1 suppression leads to mESC neuronal differentiation^17^. *PBX1* itself is also an ASD risk gene^8,21,22^. In our ASD models, we observed that PBX1 expression, particularly the PBX1a isoform, was reduced in NPCs with CTNNB1, PTEN, and DVL3 mutations, which corresponded with an increase in NPC proliferation. Notably, overexpression of PBX1a successfully rescued these proliferation abnormalities, highlighting PBX1 as a convergent target of a subset of ASD-associated mutations involved in NPC regulation.

Our findings underscore the potential role of PBX1 as a shared pathway in ASD and related neurodevelopmental disorders, providing insight into a common mechanism underlying increased NPC proliferation in ASD.

## Results

### Increased NPC prolifereation in CTNNB1, DVL3 and PTEN mutant NPCs convergence on commonly downregulated genes

We previously reported that NPCs derived from eight individuals with idiopathic autism and early brain growth exhibited increased proliferation compared to NPCs from five matched healthy controls^6^. The increased NPC proliferation is also supported in additional studies^7^, suggest increased NPC proliferation is a surrogate phenotype in a subset of ASD individuals with early brain overgrowth. We further identified that ASD linked PTEN p.I135L mutation contributed to the increased NPC proliferation using 2D NPCs derived from isogenic PTEN iPSC lines in both control and ASD genetic backgrounds^4^. To identify disease convergence related to the increased NPC proliferation, here, we focus on two additional ASD risk genes, CTNNB1 and DVL3 that are key compoents of the WNT pathway^23^. Mice carrying a stabilized version of beta-catenin (encoded by Ctnnb1) exhibited increased brain size by controling the cell cycle exit of neural precursor cells, suggesting that beta-catenin plays a role in regulating mouse brain development^24^. However, the majority of CTNNB1 variants identified in the ASD individuals are stop-gain mutations^25^, and our own work identified a stop-gain mutation in CTNNB1 that was found in an ASD individual with early brain overgrowth^6^. The impact of such CTNNB1 loss of function mutations on NPC proliferation in the human genetic background is understudied. Dishevelled (DVL) is a crucial component of the WNT signaling pathway, including the canonical pathway that regulates beta-catenin’s role in transcriptional activity^26^. We previously discovered embryonic transient cortical plate enlargement and accelerated NPC proliferation in *Dishevelled* mutant (*Dvl1*^-/-^, *Dvl3*^+/-^) mice^27^. Clinically, variants in *DVL3* are reported in individuals with ASD^8,9,28^. We chose to model heterzygous CTNNB1 p.Q76* mutation^6^ and the DVL3 p.Tyr540LeufsTer42 mutation^9^ (later refered to as DVL3^+/-^) identified from ASD individuals by creating isogenic mutant iPSCs using CRISPR/Cas9n genome editing in the same control genetic background, both unedited and edited iPSC lines were confirmed by amplicon sequencing (Extended Data Fig. 1a, 1b). Taking into account the previously generated PTEN p.WT/I135L on the same control genetic background^4^, our experimental design of modeling variants of three different ASD risk genes separately on the same control genetic background (Figure 1a) allowed us to identify potential convergent mechanisms shared by the three ASD genetic variants.

**Fig. 1.**
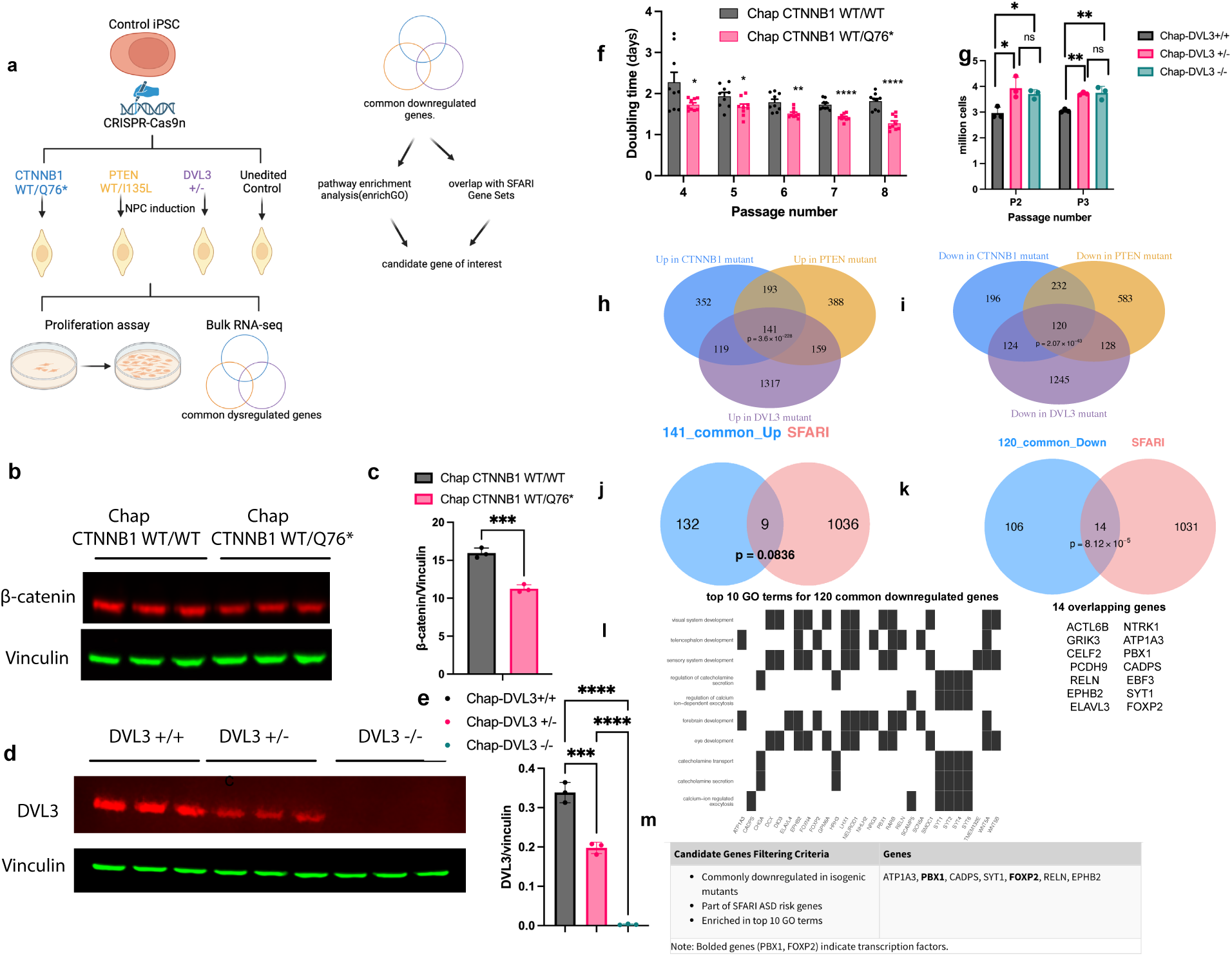
Increased NPC prolifereation in CTNNB1, DVL3 and PTEN mutant NPCs convergence on commonly downregulated genes. **a,** Workflow for characterization of isogenic iPSC-derived NPCs related to CTNNB1, PTEN and DVL3 mutations. **b,** Western blots probing for β-catenin and Vinculin from protein lysates of Chap CTNNB1 WT/WT and Chap CTNNB1 WT/Q76* at NPC passage 4. **c,** Quantification of Western blot in (b). ***p < 0.001; unpaired two-tailed t test; error bars represent SD. **d,** Western blots probing for DVL3 and Vinculin in protein lysates from Chap *DVL3*^+/+^, *DVL3*^+/-^ and *DVL3*^-/-^ at NPC passage 3. **e,** Quantification of Western blot in (d). P values were calculated with one-way ANOVA with Tukey’s correction for multiple comparisons. ***p < 0.001; ****p < 0.0001; error bars represent SD. **f,** Population doubling time assay for iPSC-derived isogenic *CTNNB1* 2D NPCs from passage 4 to passage 8. *p < 0.05; **p < 0.01; ****p < 0.0001; unpaired two-tailed *t* test; error bars represent SD. **g,** Cell proliferation quantification for isogenic *DVL3* mutant NPCs at passage 2. P values were calculated with one-way ANOVA with Holm Sidak’s correction for multiple comparisons. *p < 0.05; ns, not significant; error bars represent SD. **h,** Venn diagram illustrating the overlap of upregulated genes across 805 genes in CTNNB1 mutant NPCs, 881 genes in PTEN mutant NPCs, and 1736 genes in DVL3+/- mutant NPCs. **i,** Venn diagram showing the overlap of downregulated genes across 672 genes in CTNNB1 mutant NPCs, 1063 genes in PTEN mutant NPCs, and 1617 genes in DVL3^+/-^ mutant NPCs. **j,** Venn diagram illustrating the overlap of between 141 commonly upregulated genes across isogenic CTNNB1, PTEN and DVL3 mutant NPCs and 1045 SFARI ASD risk genes. **k,** Venn diagram illustrating the overlap of between 120 commonly downregulated genes across isogenic CTNNB1, PTEN and DVL3 mutant NPCs and 1045 SFARI ASD risk genes. The statistical significance of the overlaps of panels h, i, j and k was assessed using a hypergeometric test. **l,** heatplot visualization top 10 enriched GO terms by the 120 commonly downregulated among the three isogenic mutant NPCs. **m,** table summary of genes that are affected by all three isogenic mutant NPCs at the gene expression level and the filtering criteria used.

To determine whether the CTNNB1 p.Q76* variant and DVL3^+/-^ alter NPC proliferation, we differentiated the isogenic iPSC lines into the 2D NPCs expressing NPC markers such as PAX6, SOX2, SOX1 and Nestin (Extended Data Fig.1a-d). Both variants resulted in haploinsufficiency in NPCs, leading to reduced protein expression compared to isogenic controls (Fig.1b-e). Importantly, NPCs with the CTNNB1 p.Q76* variant and NPCs with DVL3^+/-^ variant displayed increased proliferation compared to isogenic controls using the population doubling time assay (Fig.1f-g, Extended Data Fig. 1e-f). Of note, the proliferation rate was similar between DVL3^-/-^ and DVL3^+/-^, suggesting single allele KO of DVL3, as was seen in the ASD individual^9^, is sufficient to alter NPC growth.

Since we previously profiled the 2D NPCs of the isogenic PTEN p.I135L 2D NPCs using bulk RNA-seq^4^, to identify the commonly dysregulated genes among the three ASD risk genes variants, we additionally performed bulk RNA-seq on 2D NPCs derived from control iPSC, control iPSC possessing isogenic CTNNB1 p.Q76* variant and control iPSC possessing DVL3^+/-^ variant. We found 805 upregulated genes in CTNNB1 p.Q76* mutant NPCs (Fig. 1h, Supplementary Table 1, |Log2 fold change| >0.5, p.adjust <0.05), 881 upregulated genes in PTEN p.I135L mutant NPCs (Fig. 1h, Supplementary Table 2, |Log2 fold change| >0.5, p.adjust <0.05) and 1736 upregulated genes in DVL3^+/-^ mutant NPCs (Fig. 1h, Supplementary Table 3, |Log2 fold change| >0.3, p.adjust <0.05). 141 genes are commonly upregulated in all three isogenic mutant NPCs (Fig. 1h). We also found 672 downregulated genes in CTNNB1 p.Q76* mutant NPCs(Fig. 1i, Supplementary Table 1, |Log2 fold change| >0.5, p.adjust <0.05), 1063 downregulated genes in PTEN p.I135L mutant NPCs (Fig. 1i, Supplementary Table 2, |Log2 fold change| >0.5, p.adjust <0.05) and 1617 downregulated genes in DVL3^+/-^ mutant NPCs (Fig. 1i, Supplementary Table 3, |Log2 fold change| >0.3, p.adjust <0.05). 120 genes are commonly downregulated in all three isogneic mutant NPCs(Fig. 1i). We then asked whether the commonly dysregulated genes are overrepresented in the SFARI ASD risk gene list by performing hypergeometric testing. We found 141 commonly upregulated genes were not significantly enriched in the SFARI gene list (Fig. 1j, p=0.0836), whereas the 120 commonly downregulated were (Fig. 1k, p=8.12x10^-5^). We focused on analysing these 120 significantly and commonly downregulated genes. We performed GO term analysis and found that top 10 enciched GO terms are visual system development, telecelphalon development, sensory system development, regulation of catecholamine secretion, regulation of calcium ion-dependent exocytosis, forebrain development, eye development, catecholamine transport, catecholamine secretion, calcium-ion regulated exocytosis (Fig. 1l). Out of these 120 commonly downregulate genes, 14 overlapped with the SFARI risk gene list, these include *NTRK1, ATP1A3, PBX1, CADPS, EBF3, SYT1, FOXP2, ACTL6B, GRIK3, CELF2, PCDH9, RELN, EPHB2* and *ELAVL3*(Fig. 1k). We then restricted our analysis to the SFARI ASD risk genes that are commonly downregulated in three isogenic NPC mutants and at the same time enriched in the top 10 GO terms. We ended up with 7 genes, *ATP1A3, CADPS, SYT1, RELN, EPHB2, PBX1*, and *FOXP2*. Of the 7 genes in focus, *PBX1* and *FOXP2* stood out as transcription factors (Fig. 1m).

### Altered alternative splicing in isogenic CTNNB1, PTEN and DVL3 mutant NPCs

ASD disease pathology is not only linked to altered gene expression, but also associated with altered splicing patterns^29,30^. Replicate multivariate analysis of transcript splicing (rMATS) is a tool that can quantify events including skipped exon(SE), alternative 5’ splice site (A5SS), alternative 3’ splice site(A3SS), mutally exclusive exons (MXE) and retained intron (RI) from bulk RNA-seq datasets^31,32^. We then applied rMATS to our RNA-seq datasets on 2D NPCs on three isogenic mutant NPCs on the same control genetic background to determine whether there are convergent altered alternative splicing events. We considered false discovery rates (FDR) below 0.01 as significant alternative splicing events (Fig. 2a).

**Fig. 2.**
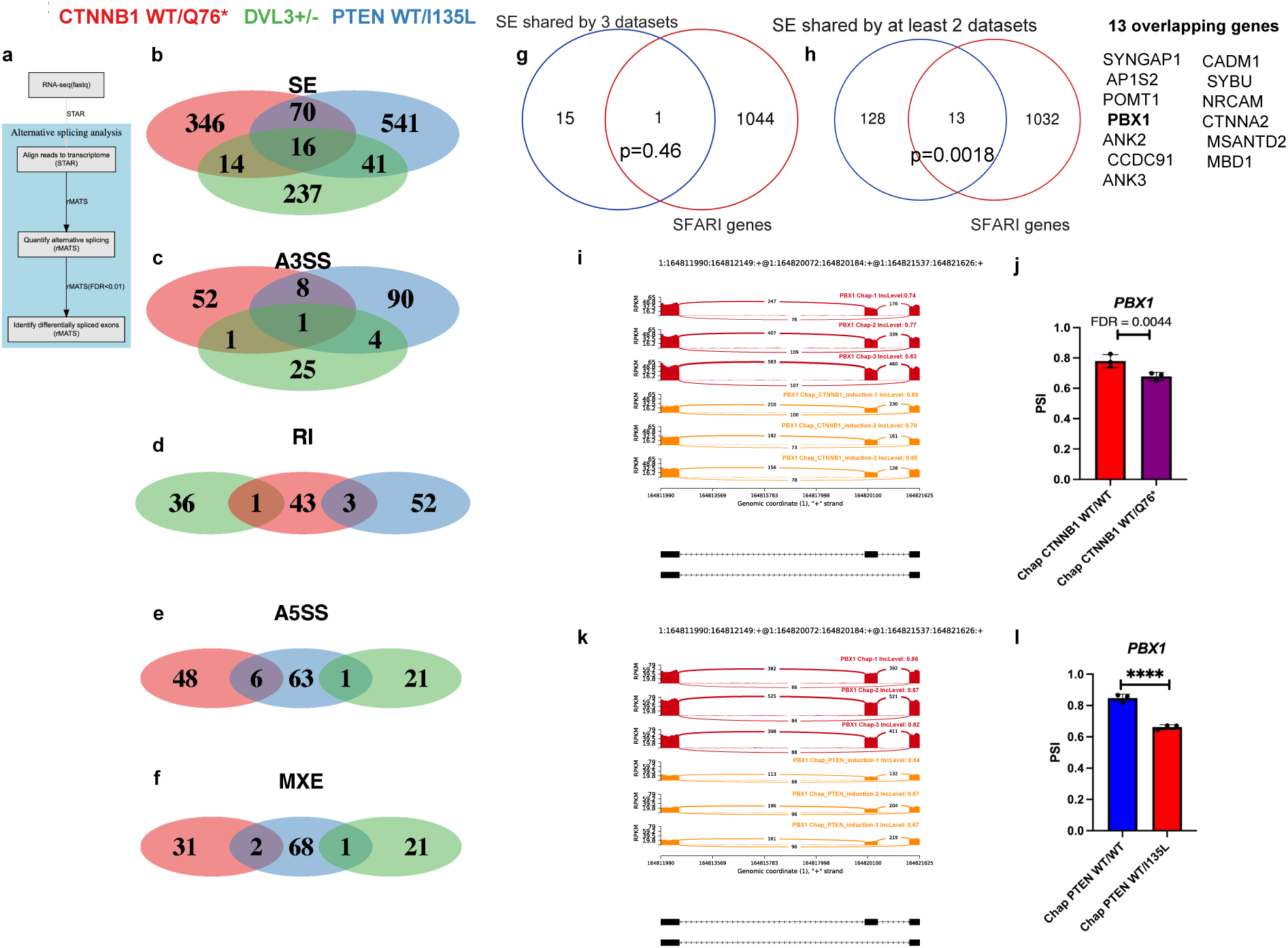
Altered alternative splicing in isogenic *CTNNB1*, *PTEN* and *DVL3* mutant NPCs. **a,** Workflow for identification of alternative splicing events in the isogenic mutant NPCs. **b,** Venn diagram for the identification of the overlapping genes among 446 genes related to SE events in CTNNB1 mutant NPCs, 668 genes related to SE events in PTEN mutant NPCs, and 308 genes with SE events in DVL3^+/-^ mutant NPCs. **c,** Venn diagram for the identification of the overlapping genes among 62 genes related to A3SS events in CTNNB1 mutant NPCs, 103 genes related to SE events in PTEN mutant NPCs, and 31 genes with SE events in DVL3^+/-^ mutant NPCs. **d,** Venn diagram for the identification of the overlapping genes among 47 genes related to RI events in CTNNB1 mutant NPCs, 55 genes related to RI events in PTEN mutant NPCs, and 37 genes with RI events in DVL3^+/-^ mutant NPCs. **e,** Venn diagram for the identification of the overlapping genes among 54 genes related to A5SS events in CTNNB1 mutant NPCs, 70 genes related to A5SS events in PTEN mutant NPCs, and 22 genes with A5SS events in DVL3^+/-^ mutant NPCs. **f,** Venn diagram for the identification of the overlapping genes among 33 genes related to MXE events in CTNNB1 mutant NPCs, 71 genes related to MXE events in PTEN mutant NPCs, and 22 genes with MXE events in DVL3^+/-^ mutant NPCs. **g,** Venn diagram illustrating the overlap between 16 genes related to SE events shared among three isogenic mutant NPCs and 1045 SFARI ASD risk genes. **h,** Venn diagram illustrating the overlap between 141 genes related to SE events shared among at least two isogenic mutant NPCs and 1045 SFARI ASD risk genes. The statistical significance of the overlap in panels g and h was assessed using a hypergeometric test. **i,** Sashimi Plot visualization for the exon 7 skipping of *PBX1* in Chap CTNNB1 WT/Q76* NPCs. The Y-axis in each plot represents a modified RPKM value. hg38 Chr1:16481990-164821628 spanning PBX1 exon 5-8 was shown. Red indicates Chap CTNNB1 WT/WT NPCs, yellow indicates Chap CTNNB1 WT/Q76* NPCs. **j,** rMATS quantification of PSI (percent spliced in) level in (i). N=3 technical replicates for Chap CTNNB1 WT/WT NPCs, N=3 technical replicates for Chap CTNNB1 WT/Q76* NPCs. **k,** Sashimi Plot visualization for the exon 7 skipping of *PBX1* in Chap PTEN WT/I135L NPCs. The Y-axis in each plot represents a modified RPKM value. hg38 Chr1:16481990-164821628 spanning *PBX1* exon5-8 was shown. Red indicates Chap PTEN WT/WT NPCs, yellow indicates Chap PTEN WT/I135L NPCs. **l,** rMATS quantification of PSI (percent spliced in) level in (k). N=3 technical replicates for Chap PTEN WT/WT NPCs, N=3 technical replicates for Chap PTEN WT/I135L NPCs. False discovery rate (FDR) was calculated by rMATS based on the Benjamini-Hochberg approach; error bars represent SD.

RNA splicing analysis revealed that the CTNNB1 WT/Q76* variant led to 446 significant events related to SE (Fig. 2b), 62 significant events related to A3SS (Fig. 2c), 47 significant events related to RI (Fig. 2d), 54 significant events related to A5SS (Fig. 2e) as well as 33 significant events related to MXE (Fig. 2f, Supplementary Table 4) compared to isogenic controls. PTEN WT/I135L variant resulted in 668 significant events related to SE (Fig. 2b), 103 significant events related to A3SS (Fig. 2c), 55 significant events related to RI (Fig. 2d), 70 significant events related to A5SS (Fig. 2e) as well as 71 significant events related to MXE (Fig. 2f, Supplementary Table 4). For DVL3^+/-^ mutant NPCs, we found 308 significant events related to SE (Fig. 2b), 31 significant events related to A3SS (Fig. 2c), 37 significant events related to RI (Fig. 2d), 23 significant events related to A5SS (Fig. 2e) as well as 22 significant events related to MXE (Fig. 2f, Supplementary Table 4). We focused on SE events as the overlap of the other four events among the three isogenic NPCs were minimal (Fig. 2b-f).

Sixteen genes displayed consistent skipped exon (SE) events across all three isogenic mutant NPC lines (Fig. 2b). however, only 1 out of the 16 genes overlap with the SFARI ASD risk gene list (Fig. 2g, p=0.46). We then asked whether SE events observed in at least 2 datasets displayed significant overlap with the SFARI gene list. Indeed, we found 13 out of 141 common SE events shared by at least two isogenic NPC datasets in control genetic background in our current study displayed significant overlap with the SFARI gene list (Fig. 2h, p=0.0018). These include *SYNGAP1, AP1S2, POMT1, PBX1, ANK2, CCDC91, ANK3, CADM1, SYBU, NRCAM, CTNNA2, MSANTD2* and *MBD1* (Fig. 2h). Surprisingly, one ASD risk gene, *PBX1*, was identified as a top candidate from both gene expression analysis and alternative splicing analysis using the three isogenic NPC bulk RNA-seq datasets (Fig. 1k, Fig. 2h). Of note, the *PBX1* gene expression patterns found in NPCs with the *CTNNB1* WT/Q76* variant, NPCs with DVL3^+/-^ variant, and NPCs with the *PTEN* WT/I135L variant were similarly downregulated (Fig. 1k, Fig. 3a, 3c & 3e). With regards to alternative splicing pattern, exon 7 skipping occured in both *PTEN* and *CTNNB1* mutant NPCs compared to isogenic controls (Fig. 2i-l). This led us to focus on *PBX1*.

**Fig. 3.**
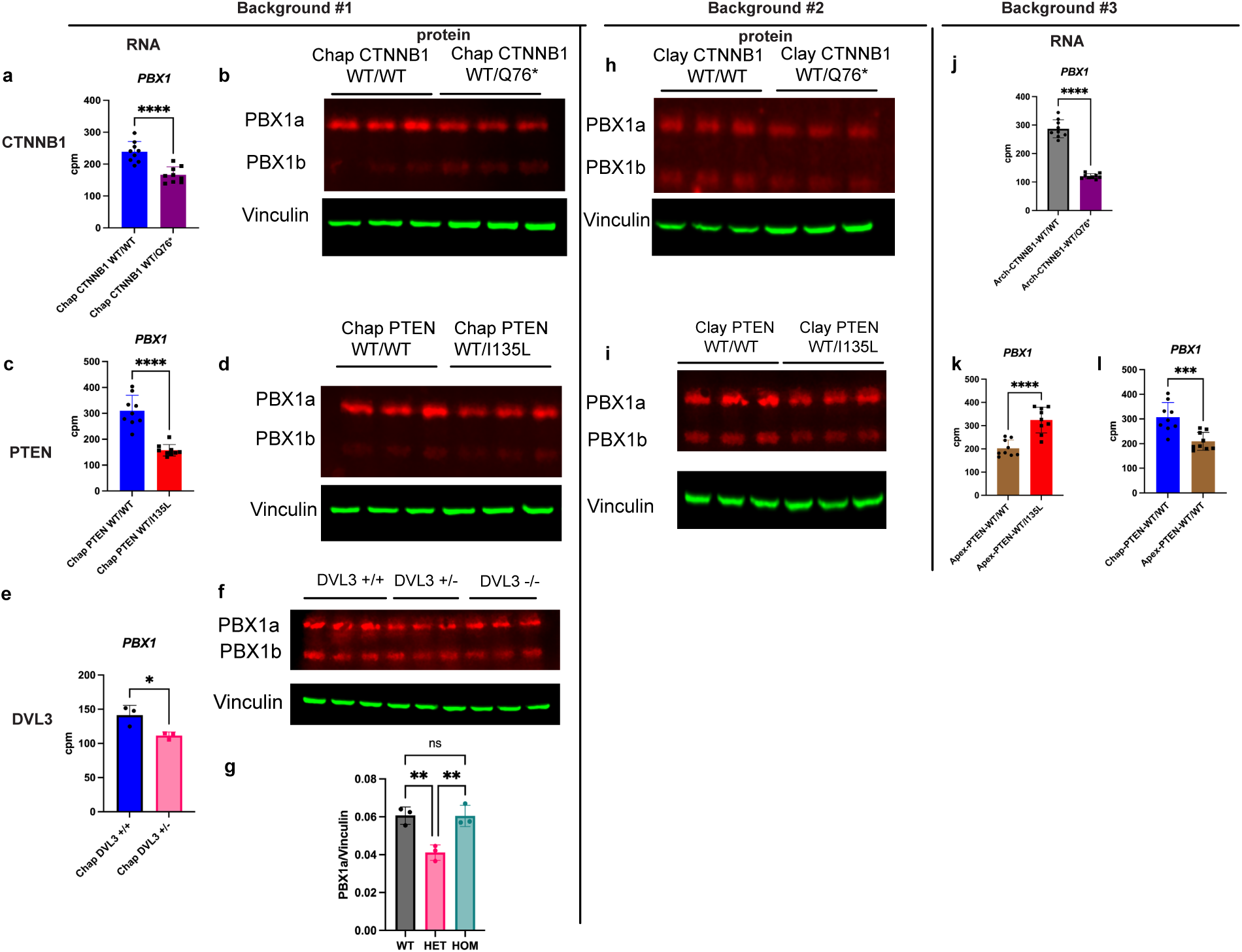
Altered PBX1 expression in isogenic CTNNB1 and isogenic PTEN mutant NPCs in multiple genetic backgrounds. **a,** RNA-seq quantification for *PBX1* in isogenic 2D CTNNB1 NPCs, demonstrates that *PBX1* RNA is decreased in *CTNNB1* mutant NPCs compared to isogenic controls. cpm, count per million. N=9 technical replicates for Chap CTNNB1 WT/WT, N=9 technical replicates for Chap CTNNB1 WT/Q76*. ****p < 0.0001; unpaired two-tailed *t* test; error bars represent SD. **b,** Western blot probing for PBX1 and Vinculin for protein lysates of Chap CTNNB1 WT/WT, Chap CTNNB1 WT/Q76* at NPC passage 4. **c,** RNA-seq quantification for *PBX1* in isogenic 2D PTEN NPCs, demonstrates that PBX1 RNA is decreased in *PTEN* mutant NPCs compared to isogenic controls. cpm, count per million. N=9 technical replicates for Chap PTEN WT/WT, N=9 technical replicates for Chap PTEN WT/I135L. ****p < 0.0001; unpaired two-tailed *t* test; error bars represent SD. **d,** Western blot probing for PBX1 and Vinculin for protein lysates of Chap PTEN WT/WT, Chap PTEN WT/I135L at NPC passage 4. **e,** RNA-seq quantification for *PBX1* in isogenic 2D *DVL3*^+/+^ and *DVL3*^+/-^ NPCs. cpm, count per million. N=3 technical replicates for Chap *DVL3*^+/+^, N=3 technical replicates for Chap *DVL3*^+/-^. *p < 0.05; unpaired two-tailed t test; error bars represent SD. **f,** Western blots probing for PBX1 and Vinculin for protein lysates of Chap *DVL3*^+/+^, *DVL3*^+/-^ and *DVL3*^-/-^ at NPC passage 3. **g,** Quantification of Western blot in (f). P values were calculated with one-way ANOVA with Tukey’s correction for multiple comparisons. **p < 0.01; ns, not significant; error bars represent SD. **h,** Western blots probing for PBX1 and Vinculin for protein lysates of Clay CTNNB1 WT/WT and Clay CTNNB1 WT/Q76* at NPC passage 2. **i,** Western blots probing for PBX1 and Vinculin for protein lysates of Clay PTEN WT/WT and Clay PTEN WT/I135L at NPC passage 2. **j,** RNA-seq quantification for *PBX1* in isogenic *CTNNB1* 2D NPCs on the Arch ASD genetic background, demonstrates that *PBX1* RNA is reduced in *CTNNB1* mutant NPCs compared to isogenic controls. cpm, count per million. N=9 technical replicates for Arch CTNNB1 WT/WT, N=9 technical replicates for Arch CTNNB1 WT/Q76*. ****p < 0.0001; unpaired two-tailed *t* test; error bars represent SD. **k,** RNA-seq quantification for *PBX1* in isogenic *PTEN* 2D NPCs on the Apex ASD genetic background, demonstrates that *PBX1* RNA is increased in *PTEN* mutant NPCs compared to isogenic controls. cpm, count per million. N=9 technical replicates for Apex PTEN WT/WT, N=9 technical replicates for Apex PTEN WT/I135L. ****p < 0.0001; unpaired two-tailed *t* test; error bars represent SD. **l,** RNA-seq quantification for *PBX1* comparing control Chap PTEN WT/WT and ASD Apex PTEN WT/WT, demonstrates that *PBX1* RNA is reduced in Apex PTEN WT/WT NPCs compared to Chap PTEN WT/WT NPCs. cpm, count per million. N=9 technical replicates for Chap PTEN WT/WT, N=9 technical replicates for Apex PTEN WT/WT. ****p < 0.0001; unpaired two-tailed *t* test; error bars represent SD.

### Altered PBX1 expression in isogenic CTNNB1 and isogenic PTEN mutant NPCs in multiple genetic backgrounds

We then determined whether PBX1 protein expression was also dysregulated in the isogenic NPC with heterozygous PTEN p.I135L variant, isogenic NPC with heterozygous CTNNB1 p.Q76* variant and isogenic NPC with DVL3^+/-^ variant by western blot analysis. We observed decreased PBX1a isoform expression in all three isogenic mutant NPCs in the control Chap genetic background (Fig. 3b, 3d, 3f-g, Extended Data Fig.1g). PBX1a was specifically decreased in DVL3^+/-^ NPCs, but remained unaltered in the DVL3^-/-^ NPCs (Fig. 2f-g), suggesting that while both homo and heterozygous DVL3 KO led to equal level of increased NPC proliferation, post transcriptional regulation may differ between DVL3^+/-^ and DVL3^-/-^ NPCs.

For CTNNB1 WT/Q76* variant, we also generated isogenic CTNNB1 mutant iPSC on a second control (Clay^6^) genetic background (Extended Data Fig.2a), and found consistent decreased PBX1a protein expression in CTNNB1 mutant NPCs (Fig. 3h, Extended data Fig.2b, 2c). Beta-catenin protein expression was also reduced (Extended data Fig.2d, 2e). In addition to the two control genetic backgrounds for CTNNB1, we also have produced isogenic CTNNB1 WT/WT and CTNNB1 WT/Q76* iPSC lines on the ASD genetic background (Arch ASD iPSC line^5,6^) (Extended Data Fig.3a). RNA-seq analysis of NPCs derived from Arch CTNNB1 WT/WT and Arch CTNNB1 WT/Q76* iPSCs (Extended Data Fig. 3b) revealed reduced *PBX1* expression in isogenic CTNNB1 WT/Q76* ASD NPCs (Fig. 3j) and identical skipped exon event in PBX1 (Fig.2j, Extended Data Fig. 3c). Thus, NPCs with CTNNB1 WT/Q76* variant displays consistent decreased PBX1 gene or protein expression in three independent genetic backgrounds.

For PTEN WT/I135L variant, we previously created isogenic *PTEN* mutant iPSCs in the control Clay genetic background^4^, so we produced 2D NPCs (Extended Data Fig.2b) and found that the effect of PTEN p.I135L variant on decreasing PBX1a protein expression in the NPC stage was replicated in this second control genetic background (Fig. 3i, Extended Data Fig.2f), accompanied by increased NPC proliferation based on a cell doubling time assay (Extended Data Fig.2g-h). We also previously derived NPCs from isogenic PTEN WT/WT and isogenic PTEN WT/I135L iPSC lines on the ASD genetic background (Apex ASD iPSC)^4^ (Extended Data Fig.3c) and performed bulk RNA-seq on the 2D NPCs^4^. The effect of the PTEN WT/I135L on *PBX1* gene expression was then examined on this third genetic background. Unexpectedly, we found the *PBX1* gene expression upregulated in the ASD PTEN WT/I135L NPCs compared to ASD PTEN WT/WT NPCs (Fig. 3k). Such oppsite direction of PBX1 regulation might be due to the effect of the strong ASD genetic background in the Apex cell line, which is consistent with our previous gene set enrich analysis (GSEA) on the effect of PTEN p.I135L variant on the control genetic background and its effect on the ASD genetic background at the NPC stage: genes related to GO terms such as regulation of neurogenesis was downregulated in the control genetic bakcgound due to PTEN p.I135L, whereas genes related to such GO terms switched to be upregulated in the ASD genetic background^4^. This ASD genetic background effect also involved dysregulated cortical neurogenesis^4^. Therefore, we then asked whether the ASD genetic background itself altered the PBX1 gene expression by comparing Apex PTEN WT/WT vs Chap PTEN WT/WT. Indeed, we found decreased *PBX1* expression in the ASD Apex PTEN WT/WT NPCs compared to control Chap PTEN WT/WT NPCs(Fig. 3i). Further analysis on *PBX1* alternative splicing revealed two independent ASD genetic backgrounds (Apex PTEN WT/WT and Arch CTNNB1 WT/WT) resulted in skipped exon (SE) events in *PBX1* (Extended Data Fig.4a-d). The *PBX1* upregulation observed in the PTEN WT/I135L mutant in the ASD genetic background is likely due to the strong effect of the ASD genetic background, which by itself alters *PBX1* gene expression.

In summary, we have identified variants in three different ASD risk genes, *CTNNB1*, *PTEN* and *DVL3*, all of which downregulated PBX1a expression in the control genetic backgrounds and at the same time increased the NPC proliferation.

### PBX1a overexpression rescues the increased NPC proliferation abnormalities

We next determined if increasing PBX1 protein expression is sufficient to rescue the increased NPC proliferation found in NPCs containing the *CTNNB1, PTEN* and *DVL3* heterozygous variants found in the ASD individuals. PBX1a is the predominant isoform expressed in 2D NPCs compared to PBX1b and is consistently downregulated in NPCs with the three ASD variants in control genetic backgrounds. To investigate its role, we stably overexpressed PBX1a in each of the three isogenic iPSC lines in the Chap control genetic background, harboring the CTNNB1 p.Q76* variant, the PTEN p.I135L variant, or the DVL3 p.Tyr540LeufsTer42. PBX1a, driven by the CAG promoter, was placed into the H11 safe harbor locus^33,34^ to avoid unintentionally perturbing genes important for cortical development. We confirmed the reduction of PBX1a protein expression in each of the three original isogenic mutant NPCs (Fig. 4a-c, Extended Data Fig.1d, Extended Data Fig.5, Extended Data Fig.6a-e), and successful overproduction of PBX1a transgene in each of the heterozygous variants by western blot (Fig. 4a-c, Extended Data Fig.6a-e). We then seeded NPCs in equal cell number for *CTNNB1* WT, *CTNNB1* heterozygous mutant and *CTNNB1* heterozygous mutant overexpressing PBX1a and performed cell counting for the NPCs at day 3. Similar experiments were also carried out for NPCs of *PTEN* WT, *PTEN* heterozygous mutant and *PTEN* heterozygous mutant overexpressing PBX1a as well as for NPCs of *DVL3* WT, *DVL3^+/-^*heterozygous mutant and *DVL3^+/-^* heterozygous mutant overexpressing PBX1a. For each of these experiments, we observed increased NPC proliferation in the isogenic mutant NPCs compared to isogenic controls, and rescue of the increased NPC proliferation in the isogenic mutant NPC overexpressing PBX1a to levels of proliferation similar to the isogenic wildtype control NPCs (Fig. 4d-f, Extended Data Fig.6f-h, Extended Data Fig.7a-c). In summary, increasing the reduced protein expression of PBX1a found in NPCs with mutations in each of three ASD risk genes that contribute to early brain overgrowth and the NPC hyperproliferation phenotype, *CTNNB1*, *PTEN* and *DVL3*, was sufficient to rescue the increased NPC proliferation.

**Fig. 4.**
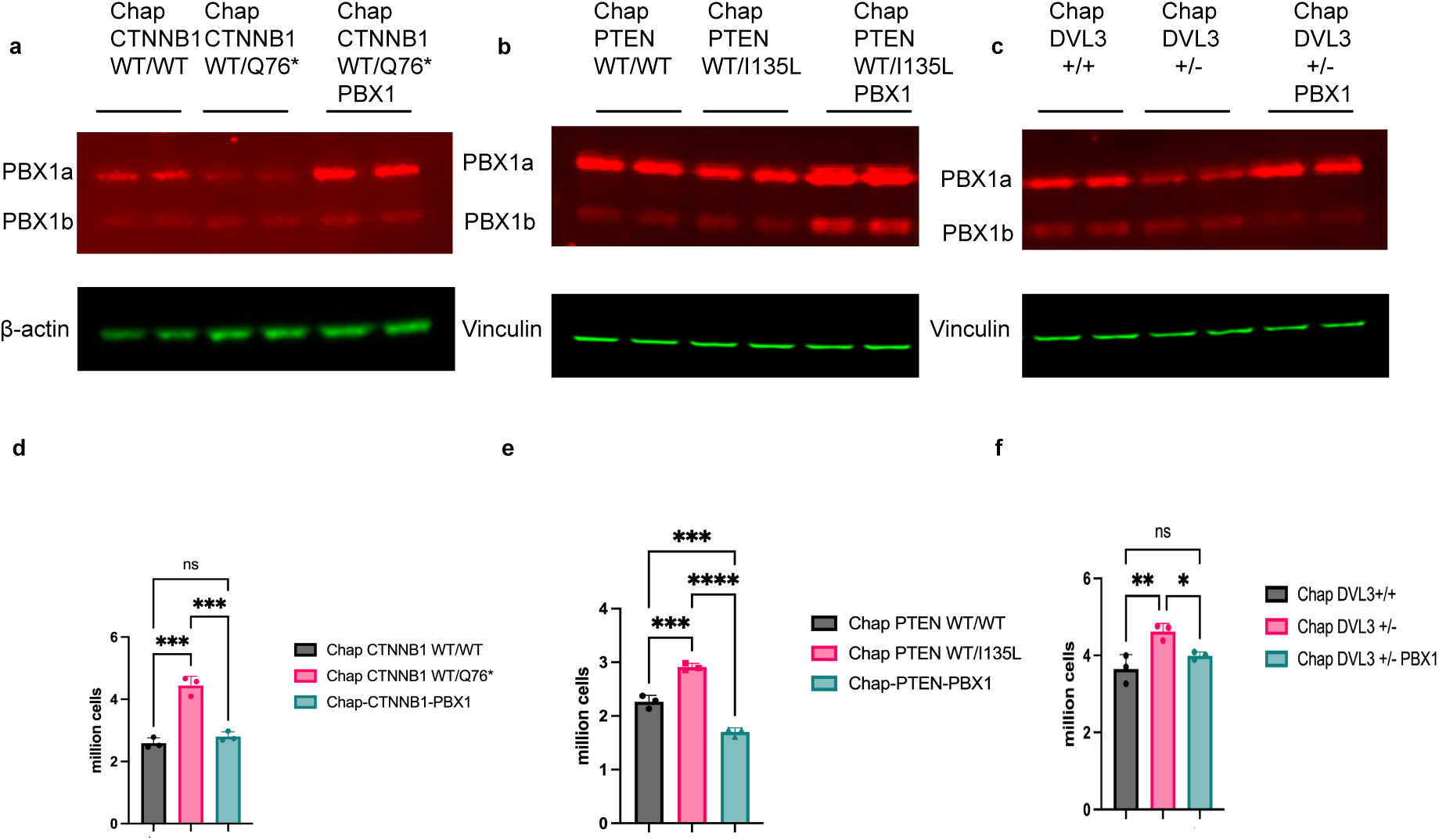
PBX1a overexpression rescues the increased NPC proliferation in 2D NPCs. **a,** Representative western blot probing for PBX1 and β-actin for protein lysates of Chap CTNNB1 WT/WT, Chap CTNNB1 WT/Q76* and Chap CTNNB1 WT/Q76* overexpressing PBX1a at NPC passage 5. **b,** Representative Western blot probing for PBX1 and Vinculin for protein lysates of Chap PTEN WT/WT, Chap PTEN WT/I135L and Chap PTEN WT/I135L overexpressing PBX1a at NPC passage 4. **c,** Representative Western blot probing for PBX1 and Vinculin for protein lysates of Chap *DVL3*^+/+^, Chap *DVL3*^+/-^ and Chap *DVL3*^+/-^ overexpressing PBX1a at NPC passage 3. **d,** Cell proliferation quantification for Chap CTNNB1 WT/WT, Chap CTNNB1 WT/Q76* and Chap CTNNB1 WT/Q76* overexpressing PBX1a at NPC passage 5. P values were calculated with one-way ANOVA with Sidak’s correction for multiple comparisons. ***p < 0.001; ns, not significant; error bars represent SD. **e,** Cell proliferation quantification for Chap PTEN WT/WT, Chap PTEN WT/I135L and Chap PTEN WT/I135L overexpressing PBX1a at NPC passage 5. P values were calculated with one-way ANOVA with Sidak’s correction for multiple comparisons. ***p < 0.001; ****p < 0.0001; error bars represent SD. **f,** Cell proliferation quantification for Chap *DVL3*^+/+^, Chap *DVL3*^+/-^ and Chap *DVL3*^+/-^ overexpressing PBX1a at NPC passage 3. P values were calculated with one-way ANOVA with Sidak’s correction for multiple comparisons. *p < 0.05; **p < 0.01; ns, not significant; error bars represent SD.

### PBX1 dysregulation identified from additional iPSC models of ASD

We have shown that increased proliferation in NPCs containing either of two ASD risk genes related to the WNT pathway or a third ASD risk gene, PTEN, converge on altering PBX1 expression. To determine whether a similar alteration in *PBX1* expression in NPCs can be found in other ASD models, we focused on publicly available RNA-seq datasets from isogenic iPSC models, aiming to uncover potential dysregulation of *PBX1*. We found that several iPSC models of ASD risk genes implicated in macrocephaly displayed *PBX1* alterations (4 out of 6 ASD risk genes, 66.7%, Supplemental Table 5). For example, NPC models of *TSC1* KO^35^ displayed downregulated *PBX1* gene expression (Extended Data Fig. 8a). *CHD8* heterzygous KO cerebral organoids^36^ also exhibited decreased *PBX1* gene expression (Extended Data Fig. 8b). Loss of *FMR1* in the NPCs^37^ resulted in *PBX1* downregulation (Extended Data Fig. 8c). *PBX1* downregulation was also seen in NPCs with heterozygous *NTRK2* KO^38^ (Extended Data Fig. 8d). We analyzed RNA-seq datasets from isogenic NPC models of ASD risk genes in individuals with microcephaly, identifying *PBX1* dysregulation in 3 out of 10 ASD risk genes (30%, Supplemental Table 5) associated with mutations in *HNRNPU*, *TBL1XR1*, and *SETBP1*. Specifically, downregulated *PBX1* expression was observed in human iPSC models with heterozygous *HNRNPU* KO^39^ (Extended Data Fig. 8e), while upregulated *PBX1* expression was found in mouse cortical NPCs with *Tbl1xr1* KO^40^ (Extended Data Fig. 8f) and in human iPSC-derived NPCs with *SETBP1* KO^41^ (Extended Data Fig. 8g). Among iPSC-derived NPC models with mutations in ASD risk genes not associated with early brain overgrowth, we identified two genes that exhibited *PBX1* downregulation when mutated (2 out of 5 ASD risk genes, 40%, Supplemental Table 5): *DLG2*^42^ (Extended Data Fig. 8h) and *POU3F2*^43^ (Extended Data Fig. 8i).

Upon querying the previously published bulk RNA-seq analysis of ASD vs control prefrontal cortex (PFC)^44^, we found that *PBX1* downregulation was found in the PFC of ASD individuals (Extended Data Fig. 8j), suggesting in addition to NPC, neurons and/or glia cells are likely to be cell types that display *PBX1* dysregulation in individuals with ASD.

In summary, these additional bulk RNA-seq dataset analysis provide evidence that the *PBX1* dysregulation is not limited to the three ASD risk genes in our own study. Mutations in other ASD risk genes related to macrocephaly, microcephaly or ASD risk genes unrelated to brain size may also lead to altered *PBX1* gene expression in NPCs, an interpretation that is further strengthened with the observed *PBX1* downregulation in the PFC of the ASD Postmortem brains.

## Discussion

Uncovering convergent mechanisms in ASD pathology has proven challenging^45–48^, largely due to the inherent genetic and phenotypic heterogeneity of ASD and the scarcity of studies that model individual ASD-linked gene mutations in isogenic iPSC lines to explore disease convergence^2^. In this study, we generated isogenic iPSC lines for three ASD risk genes and profiled the transcriptomes of iPSC-derived NPCs, with a focus on gene expression and alternative splicing to identify key candidate genes. Our results highlight a convergence among these ASD-linked mutations, leading to enhanced NPC proliferation alongside reduced expression of PBX1 mRNA and diminished levels of its PBX1a protein isoform. Importantly, we successfully rescued heightened NPC proliferation in isogenic mutant NPCs for each of the three ASD risk genes, *CTNNB1*, *PTEN*, and *DVL3*, by stably overexpressing PBX1a. We further confirmed *PBX1* alternation using iPSC-derived NPCs from additional genetic backgrounds with the same *PTEN* and *CTNNB1* mutations.

Our findings suggest that *PBX1* serves as a convergent target of some ASD-associated mutations, linking mutations in *CTNNB1*, *PTEN*, and *DVL3* to increased NPC proliferation, a surrogate marker for brain overgrowth during neurogenesis. PBX1a, the primary isoform affected, plays a critical role in mouse embryonic cortex development, including dorsal-ventral patterning and neural differentiation^18^, while PBX1b is involved in rodent adult neurogenesis within other brain regions such as the SVZ^19^. This isoform-specific activity highlights PBX1’s dual role in developmental and adult neurogenesis. Our data suggest that PBX1a downregulation in NPCs may drive the hyperproliferative phenotype observed in human NPC models of ASD. In the mouse, the splicing factor Ptbp1 suppresses Pbx1a isoform expression by skipping exon 7 in mouse embryonic stem cells (mESCs); removal of this suppression by Ptbp1 leads to mESC neuronal differentiation. PBX1 itself is also recognized as an ASD risk gene, underscoring its relevance in neurodevelopmental pathways. In addition, we uncovered similar *PBX1* dysregulation by mutations in other ASD-linked genes such as *TSC1, CHD8, FMR1, HNRNPU, DLG2, NTRK2, POU3F2, TBL1XR1* and *SETBP1* using published datasets. Bulk RNA-seq analysis of the PFC in ASD compared to controls revealed downregulation of *PBX1*^44^. Together, these findings underscore *PBX1* as a crucial target identified across both in vitro stem cell models of ASD and ASD patient brain samples.

An important role for *PBX1* dysregulation has been established in several types of cancers^13–16^. PBX1 regulates proliferation in human cancers dependent upon context: PBX1 downregulation inhibited breast cancer and small cell lung cancer growth, while reduced PBX1 increased colorectal cancer proliferation^49^. Mouse studies have documented a critical role for *Pbx1* during neurogenesis in the adult subventricular zone^19^ and olfactory bulb^20^, though not during embryonic neurogenesis. Here we demonstrate that *PBX1* dysregulation can also be observed in developmental neurological disorders. *PBX1* is a cofactor with Hox/HOM-C proteins that regulate downstream target gene expression in leukemogenesis^50^. Additionally, PBX1 interacts with other homeodomain containing transcription factors that are part of the ENGRAILED family^51^, providing a potential pathway to regulate neurogenesis. Future work to determine relevant interaction partners of *PBX1* in NPC and neurons represents an exciting avenue to further understand *PBX1* dysregulation in neurological and neurodevelopmental disorders.

*PBX1* has been implicated in various developmental processes, with mutations known to cause congenital abnormalities such as congenital abnormalities of the kidney and urinary tract(CAKUT), intellectual disability, and, in some cases, macrocephaly^52,53^. However, in this study, we did not examine *PBX1* mutations directly. Instead, our data suggest that mutations in several ASD risk genes—particularly those associated with macrocephaly—converge on *PBX1* dysregulation. This convergence highlights *PBX1*’s potential role as a shared downstream target that links diverse ASD-associated mutations to increased NPC proliferation. The complex and varied impacts of PBX1 dysregulation in different developmental contexts, as seen in studies of CAKUT and other growth disorders, underscore *PBX1*’s relevance in regulating developmental pathways and suggest that *PBX1* could contribute to the neurodevelopmental and cellular phenotypes observed in ASD models.

Overall, our study reveals *PBX1* as a novel convergent target dysregulated by mutations in multiple ASD risk genes in NPCs, offering new insights into disease biology of ASD and other neurodevelopmental disorders.

## Methods

### iPS cell culture

Two control induced pluripotent stem cell (iPSC) lines, Chap and Clay^6^, were engineered to carry the CTNNB1 p.Q76* mutation using CRISPR/Cas9n genome editing^54^. These control lines had also been previously modified to harbor the patient-specific PTEN p.I135L mutation^4^. Additionally, the Chap iPSC line was further edited using CRISPR/Cas9n to create lines with DVL3 p.Tyr540LeufsTer42 variant, identified in a previous study^9^. This variant is referred to as DVL3^+/-^ to denote the ASD-specific heterozygous *DVL3* variant, while the homozygous mutation affecting both alleles is denoted as DVL3^−/−^.

ASD iPSC Apex, harboring the PTEN p.I135L mutation, and isogenic line with *PTEN* correction were previously generated^4,6^. Similarly, ASD iPSC Arch, carrying CTNNB1 p.Q76* mutation, and its isogenic line with *CTNNB1* correction were also previously generated^4,6^.

Stable iPSC lines overexpressing PBX1a at the H11 locus were generated in three isogenic mutant iPSC lines derived from the control Chap genetic background using TALEN-mediated genome editing^55^. The iPSC lines were cultured using a feeder-free protocol in six-well plates coated with Vitronectin (Gibco, A31804) and maintained in mTeSR plus medium (StemCell Technologies). The cultures were incubated at 37℃with 5% CO2, and cells were fed every other day with 2 mL of mTeSR plus medium per well. For passaging, iPSC colonies were dissociated using 0.5 mM EDTA.

### CRIPSR-Cas9 nickase mediated genome editing

We modified the original Addgene plasmid AIO-Puro (#74630)^56^ by replacing the CMV promoter with the CAG promoter and referred to the resulting construct as CAG-AIO-puro. For the *CTNNB1* mutant, specific guide RNAs (gRNAs), with sequences 5’gttcccactcatacaggact 3’ and 5’ ttcactcaagaacaagtagc 3’, were individually integrated into the CAG-AIO-puro plasmid. The donor plasmid was crafted by amplifying 800 bp segments upstream and downstream of the mutant site using genomic DNA from the ASD iPSC line Arch, bearing the CTNNB1 p.Q76* mutation. The resulting PCR product underwent KpnI and BamHI digestion before being cloned into the pUC19 plasmid. Sanger sequencing confirmed the successful construction of the donor plasmid designed to introduce the CTNNB1 p.Q76* mutation into the *CTNNB1* locus.

Co-transfection of the gRNAs and donor plasmid into the control iPSC line Chap^6^ was facilitated by Lipofectamine™ Stem Transfection Reagent. Puromycin was administered from day 1 to day 3. Subsequently, iPSCs were dissociated into single cells using TrypLE and seeded at a density of 200 cells per well in a 6-well plate. Single colonies were then selected, expanded and subjected to Sanger sequencing followed by Miseq analysis to verify the generation of genome-edited iPSC clones.

For *DVL3* mutant iPSC lines, guide RNAs with sequences 5’tactggtacgggaaagccat 3’ and 5’ gcacccatacaacccgcacc 3’ were introduced into the CAG-AIO-puro plasmid. A 637 bp PCR product, comprising 330 bp upstream and 308bp downstream of the intended targeting site, was subsequently amplified. The c.1618T duplication was incorporated using site-directed mutagenesis via the Q5 Site-Directed Mutagenesis Kit.

For genotyping convenience, both *CTNNB1* and *DVL3* donor plasmids included a 22bp random sequence ‘ccgcagaaatacattgaatcgg’ downstream of the mutation site in the intron region, achieved through the Q5 Site-Directed Mutagenesis Kit. Sanger sequencing was conducted to confirm the successful generation of the *DVL3* donor plasmid. Transfection of gRNAs and donor plasmids for *DVL3* was carried out similarly to *CTNNB1* genome editing, with confirmation of genome-edited iPSC clones accomplished through targeted Miseq analysis.

### TALEN mediated genome editing for PBX1a overexpression

We digested plasmid H11-CAG-roxSTOProx-Zsgreen-NLS^5^ with NruI and EcoRI. Subsequently, gel extraction was performed to eliminate roxStoprox_Zsgreen, retaining the upper band as the vector for subsequent cloning steps. cDNA was then synthesized from 600ng of RNA obtained from the control Chap NPC using Superscript III First-Strand Synthesis System (Thermo Fisher). The PBX1a isoform was PCR amplified using Platinum™ SuperFi II PCR Master Mix (Thermo Fisher) and cloned into the prepared vector through NEBuilder HiFi DNA Assembly (New England Biolabs). The resulting donor plasmid, named H11-CAG-PBX1a, for overexpressing PBX1a in the H11 safe harbor locus, was verified through Sanger sequencing.

For transfection into individual isogenic Chap *PTEN*, *CTNNB1*, and *DVL3* mutant iPSC lines, addgene plasmids #51554 and #51555 (50 ng each) were co-transfected with 400 ng of the donor plasmid in one well of a 24-well plate. G418 was utilized to select for edited clones over a week. Following this, cells were dissociated into single cells and seeded at a low density (200 cells per well in a 6-well plate) to facilitate the formation of separated colonies. PCR genotyping, employing the primers 5’ H11-screening-F TCTGACCCTTTAGCATGGAGTC and 5’ H11-screening-R GCTTCGGTGTGTCCGTCA with Gotaq master-mix, was conducted. Successful overexpression of PBX1a was further confirmed at the NPC stage through western blot analysis for all the edited cell lines.

### Confirmation of edits using Miseq

For isogenic *CTNNB1*, *PTEN* and *DVL3* mutants, we prepared PCR amplicon near the targeting region, and performed Miseq based next generation sequencing to confirm the edits and no unintended indels produced using automatic python package CRISPResso^57^ (version: 2.2.7).

### Dorsal NPC production from iPS cell lines

For data presented related to isogenic CTNNB1 panel (Chap CTNNB1 WT/WT, Chap CTNNB1 WT/Q76*), isogenic PTEN panel (Chap PTEN WT/WT, Chap PTEN WT/Q76*) on the chap control genetic background, Isogenic CTNNB1 panel (Arch CTNNB1 WT/WT, Arch CTNNB1 WT/Q76*) and isogenic PTEN panel (Apex PTEN WT/WT, Apex PTEN WT/I135L) on the ASD genetic background, a commercial kit from Thermo Fisher was used as previously described^4^. In brief, on day (-1), iPSCs were dissociated with EDTA and seeded at approximately 25% confluence in mTeSR plus media in vitronectin coated plate, on day 0, media was replaced with 2 ml NIM medium, medium change was performed on day 2, 4, 5 and 6. On day 7, cells were dissociated using accutase and seeded in NEM media at 1 million cells per well in 6-well plates coated with Geltrex. We also supplemented media with 5 uM ROCK inhibitor to improve survival. On the second day, NEM media was refreshed to remove the ROCK inhibitor. Media change was then performed every other day. This is considered passage 0. Cells were then passaged using Accutase.

For all other data presented including isogenic CTNNB1 panel on the Clay control genetic background (Clay CTNNB1 WT/WT, Clay CTNNB1 WT/Q76*), isogenic PTEN panel on the Clay control genetic background (Clay PTEN WT/WT, Clay PTEN WT/I135L), isogenic DVL3 panel (Chap DVL3^+/+^, Chap DVL3^+/-^, Chap DVL3^-/-^), isogenic PBX1 rescue panel in Fig.4 (Chap CTNNB1 WT/WT, Chap CTNNB1 WT/Q76* and Chap CTNNB1 WT/Q76* overexpressing PBX1a; Chap PTEN WT/WT, Chap PTEN WT/I135L, Chap PTEN WT/I135L overexpressing PBX1a; Chap DVL3^+/+^, Chap DVL3^+/-^, Chap DVL3^+/-^ overexpressing PBX1a), we used a previously published NPC procotol^58,59^. In brief, iPSCs were dissociated and seeded in mTeSR™ Plus media in 12-well plates coated with vitronectin at approximately 25% confluence at day (-1). On day 0, media was removed, and replaced with 1 ml NPC basal media freshly supplemented with 4µM CHIR99021, 3µM SB431542 and 0.1 µM Compound-E. Media change was performed on day 2, 4, 5 and 6, and on day 7, cells were dissociated with accutase and seeded in 1 million cells per wells in Matrigel coated 6-well plate in NPC basal media freshly supplemented with 3 µM CHIR99021 and 2 µM SB431542 as well as 5 uM ROCK inhibitor. We considered cells at this point as passage 0. On the second day, media change was performed to remove the ROCK inhibitor, media change was then done every other day. NPC basal media is composed of Neurobasal: Advanced DMEM/F12 (1:1), 1xN2, 1xB27 (Catalog number: 17504044), 1% Glutmax, 5 µg/ml BSA and 10 ng/mL hLIF.

### Immunofluorescence

Immunofluorescence was performed as was previously described with modifications^4^. In brief, 2D NPCs were washed with PBS, and fixed with 4% paraformaldehyde (PFA) for 15 mins at room temperature. After three washes, cells were permeabilized for 10 min using 0.1% Triton X-100 in PBS, and blocked for at least 1 h at room temperature with blocking buffer (0.1% Triton X- and 1% BSA in PBS). Primary antibody was incubated overnight in 4°C in blocking buffer. On the second day, cells were washed three times using PBS, and then incubated with the appropriate secondary antibody at room temperature for 30 min. Cells were then counterstained with DAPI in PBS and followed by 3 washes with PBS. For Extended Data Fig. 1c-d, Extended Data Fig. 3b, 3e, images were captured using Leica DM6000 inverted microscope in the CWRU Light Microscopy Imaging Core. For Extended Data Fig.2b and Extended Data Fig.5, images were captured using EVOS FL Color Imaging Systems.

### Western blot

NPCs cultured in 6 well plates were collected once reaching 90% confluence. Briefly, media was first aspirated, then cells were washed with PBS, then cold PBS was added, and cells were scraped off the plate. Cells were collected by centrifugation at 1200 rpm for 3 min, PBS was aspirated, and cell pellets were resuspended in Mammalian Protein Expression Reagent (MPER) supplemented with Halt Protease and Phosphatase Inhibitor Single-Use Cocktail. This was followed by 25 min incubation on ice. Samples were centrifuged at 14,000 rcf for 15min, then protein lysates were aliquoted and frozen down in -80°C for future use. Protein lysates were quantified using Bradford protein assay. 8-10 ug was used for loading into the Nupage 4%-12% 17-well gel and transferred to nitrocellulose membranes. Membrane was blocked with Intercept (TBS) Blocking Buffer for 1h, then primary antibody was incubated overnight in TBS blocking buffer supplemented with 0.2% Tween-20. Blots were washed 4 times with TBS with 0.2% Tween-20, followed by incubation with fluorescent secondary antibody, then blots were washed for 4 times. Blots were then captured and quantified using the Odyssey XF Imaging System. Western blots for NGN2 induced neuron protein lysates was performed in the same manner as 2D NPCs.

### Bulk RNA-seq

#### RNA extraction

2D NPCs were first washed with DPBS, then DPBS was aspirated, 1 ml Trizol was added into 2D NPCs in one well of 6-well plates, cells were scraped off the plates and frozen in -80°C for future use. RNA was extracted using the Direct-zol RNA Miniprep Plus Kit following the kit protocol. The extracted RNA was then submitted to Novogene for library preparation and followed by paired end 150bp poly-A RNA-seq on Novaseq 6000. All samples were sequenced at approximately 30 million paired-end reads.

#### RNA-seq Data analysis

Gene expression analysis: The fastq files underwent alignment to the human reference transcriptome (Ensembl version 103) through kallisto 0.46.2. Subsequently, the resulting transcript abundance was processed following previously outlined procedures^4^. In summary, the R package Tximport was utilized for summarizing transcript quantification to genes. Normalization was then carried out using the edgeR TMM method. Genes with zero counts per million (cpm) were filtered out, and the limma voom function was applied for variance stabilization on the normalized and filtered data. Finally, differential expression analysis was conducted using limma, incorporating Benjamini-Hochberg multiple testing correction.

Splicing analysis using rMATS: To generate BAM files, we initially aligned the FASTQ files with the human reference transcriptome (Ensembl version 103) using STAR^60^ (Spliced Transcripts Alignment to a Reference) version 2.7.10a. Subsequently, replicate multivariate analysis of transcript splicing (rMATS version 4.1.2)^31^ was utilized for processing the resulting BAM files and quantifying splicing events. The Sashimi Plots were then crafted using the rmats2sashimiplot Python package, an integral part of the rMATS output analysis.

## Data and materials availability

Raw RNA-seq data for isogenic *PTEN* NPCs was deposited in Gene Expression Omnibus (GEO): GSE214323, RNA-seq data and processed files for isogenic *CTNNB1* NPCs and isogenic *DVL3* NPCs will be made available upon official publication.

This paper does not produce original code.

Any additional information required to reanalyze the data reported in this paper is available upon request.

## Acknowledgments

We thank Dr. Paul Tesar for comments on the manuscript; Drs. Fulai Jin and Yan Li for access to the EVOS FL Color Imaging Systems, Drs. Ashleigh Schaffer and Helen Miranda for access to the Zeiss Axio Observer Fluorescent Microscope and Odyssey XF Imaging System, and CWRU Light Microscopy Imaging Core for access to the Leica DM6000 inverted microscope. This study was supported by grants from the National Institutes of Health and the National Institute for Mental Health: RO1MH114601 and RO1MH113106 to AWB, and NIH Grant S10-RR021228 to CWRU SOM Light Microscopy Core Facility (Leica DM6000 inverted microscope).

## Author contributions

Conceptualization: SF.; Methodology: SF.; Investigation: SF.; Visualization: SF.; Funding acquisition: AWB.; Supervision: AWB.; Writing – original draft: SF.; Writing – review & editing: SF, AWB

## Inclusion & ethics statement

All researchers that fulfill authorship criteria have been included in the author list.

---

**Extended Data Fig. 1.**
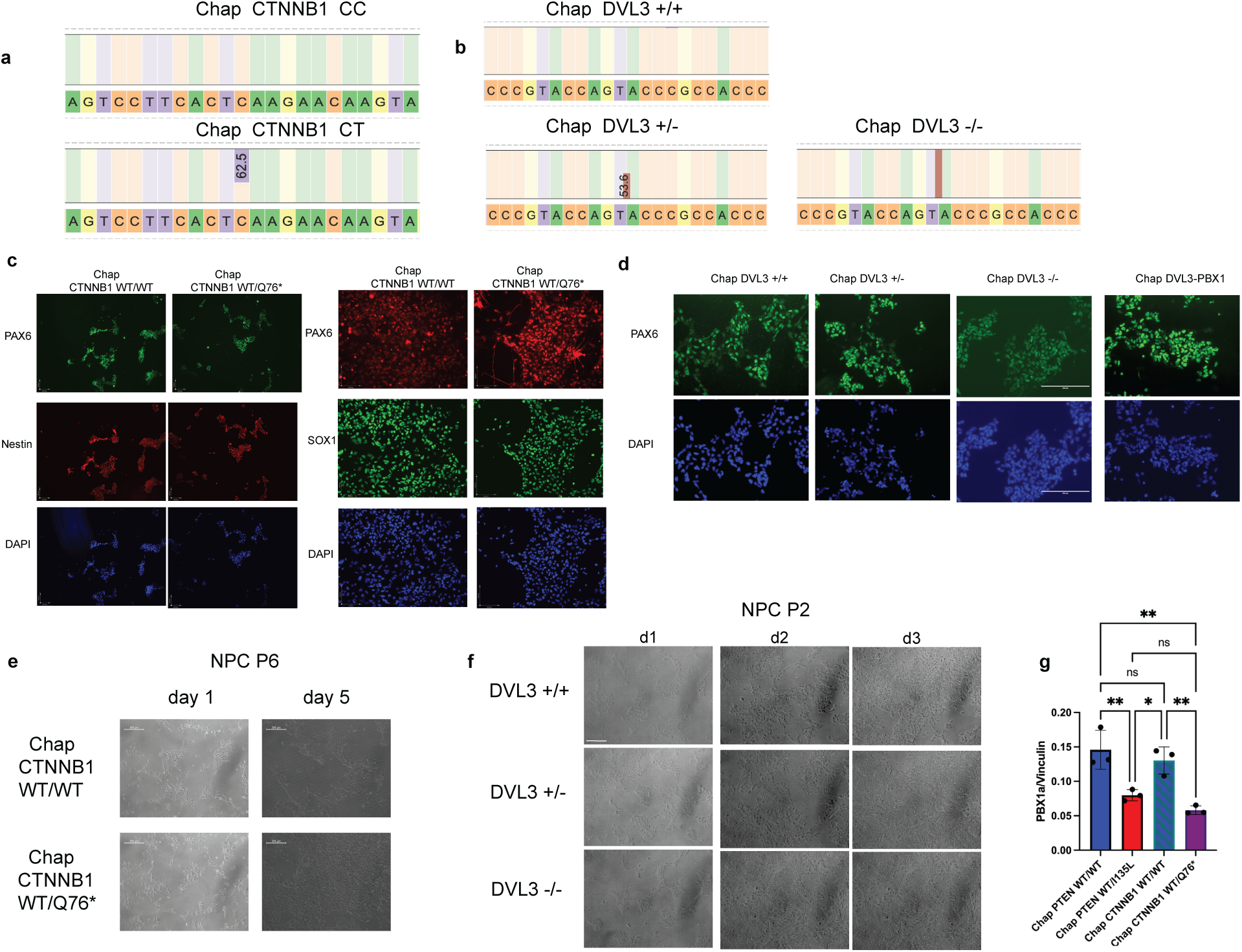
Generation and characterization of isogenic CTNNB1 p.Q76* iPSC line and isogenic PTEN p.I135L iPSC line. **a,** Mi-seq confirmation of isogenic *CTNNB1* mutant iPSC line. **b,** Mi-seq confirmation of isogenic *DVL3^+/-^* and *DVL3*^-/-^ mutant iPSC lines. Insertion indicates T duplication, resulting in DVL3 KO due to frameshift. **c,** Staining for NPC markers in NPCs derived from control iPSC (Chap CTNNB1 WT/WT) and isogenic Chap CTNNB1 WT/Q76* mutant iPSCs using the PSC Neural Induction Medium protocol, Scale bar 100µm. **d,** PAX6 staining of NPCs generated using the Li et al. 2011 protocol, including Chap DVL3+/+, Chap DVL3+/-, Chap DVL3-/-, and Chap DVL3+/- overexpressing PBX1a. Scale bar: 200 µm. **e,** Representative brightfield images for Chap CTNNB1 WT/WT and Chap CTNNB1 WT/Q76* NPCs at day1 and day5 of passage 6. Cells were seeded at equal cell numbers for both genotype. Scale bar: 200 µm. **f,** Representative brightfield images for 2D NPCs Chap DVL3 WT/WT, Chap DVL3+/- and Chap DVL3-/- at day1, day2 and day3 of passage 2. Cells were seeded at equal cell numbers for all genotypes. Scale bar: 200 µm. **g,** Quantification of PBX1a/Vinculin level in western blot of isogenic PTEN and isogenic CTNNB1 NPCs in control Chap genetic background at NPC passage 4. P values were calculated with one-way ANOVA with Tukey’s correction for multiple comparisons. *p < 0.05; **p < 0.01; ns, not significant; error bars represent SD.

**Extended Data Fig. 2.**
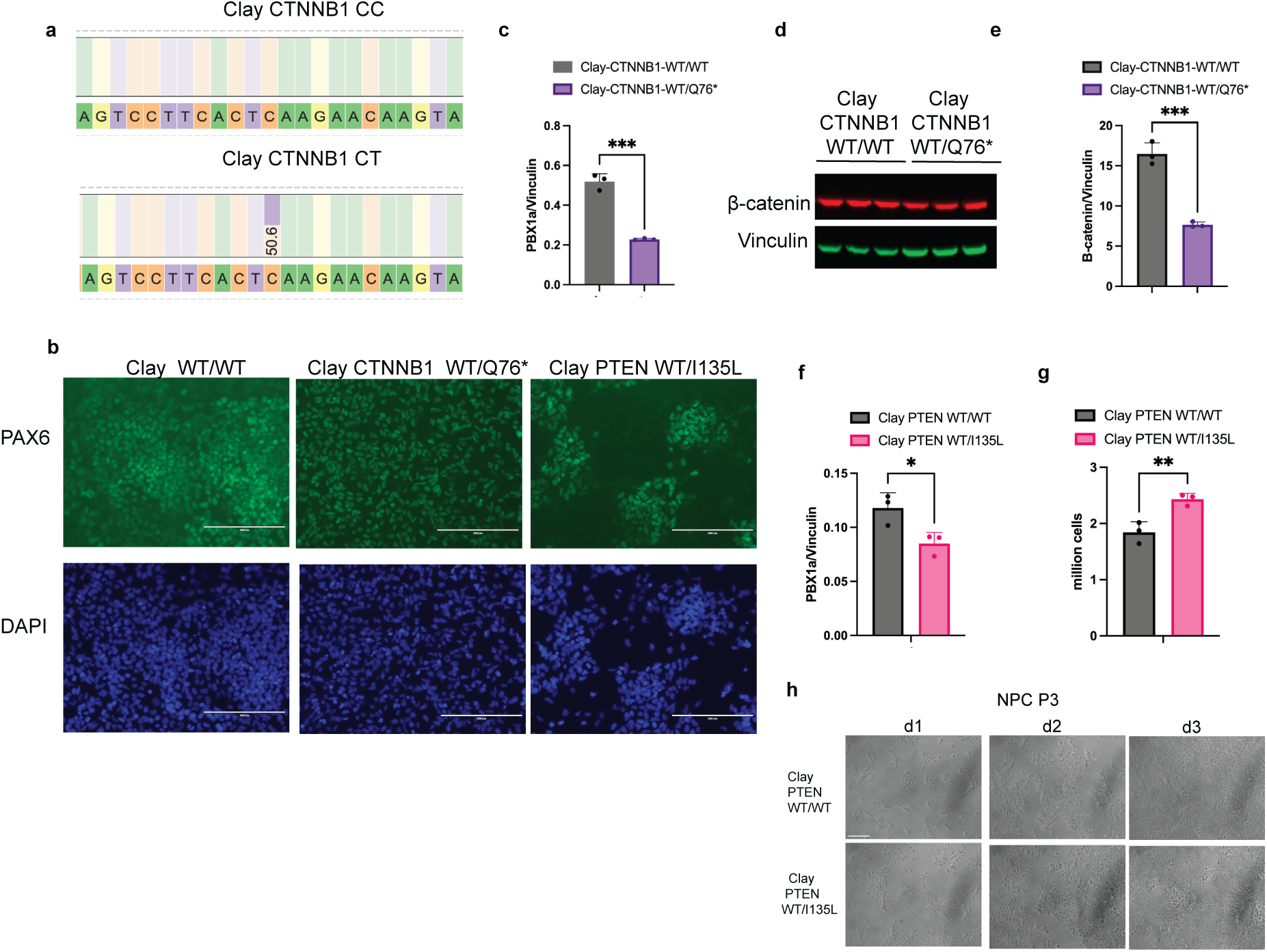
Characterization of isogenic 2D NPCs derived from PTEN p.I135L and CTNNB1 p.Q76* iPSCs on the control Clay genetic background. **a,** Mi-seq confirmation of isogenic *CTNNB1* mutant iPSC line on the control Clay genetic background. **b,** PAX6 staining for NPCs generated using the Li et al. 2011 protocol for control Clay (Clay PTEN WT/WT) and Clay with the PTEN p.Q76* mutation (Clay PTEN WT/I135L) previously installed by genome editing^4^ and Clay CTNNB1 WT/Q76* generated in this study. Scale bar 200µm. **c,** Quantification of Western blots in Fig.3b. ***p < 0.001; unpaired two-tailed t test; error bars represent SD. **d,** Western blots probing for β-catenin and Vinculin for protein lysates of Clay CTNNB1 WT/WT and Clay CTNNB1 WT/Q76* at NPC passage 2. **e,** Quantification of Western blots in (**d**). ***p < 0.001; unpaired two-tailed t test; error bars represent SD. **f,** Quantification of Western blots in Fig.3d. *p < 0.05; unpaired two-tailed t test; error bars represent SD. **g,** Cell proliferation assay for day 3 NPCs of Clay PTEN WT/WT and Clay PTEN WT/I135L. NPC Passage 3 at day 0 for both genotypes were seeded 4x10^5 cells per well in 6-well plates, and cell counting was performed on day 3; error bars represent SD. **h,** Representative brightfield images for 2D NPCs Clay PTEN WT/WT and Clay PTEN WT/I135L at day1, day2 and day3 of passage 3. Cells were seeded at equal cell numbers for both genotypes. Scale bar: 200 µm.

**Extended Data Fig. 3.**
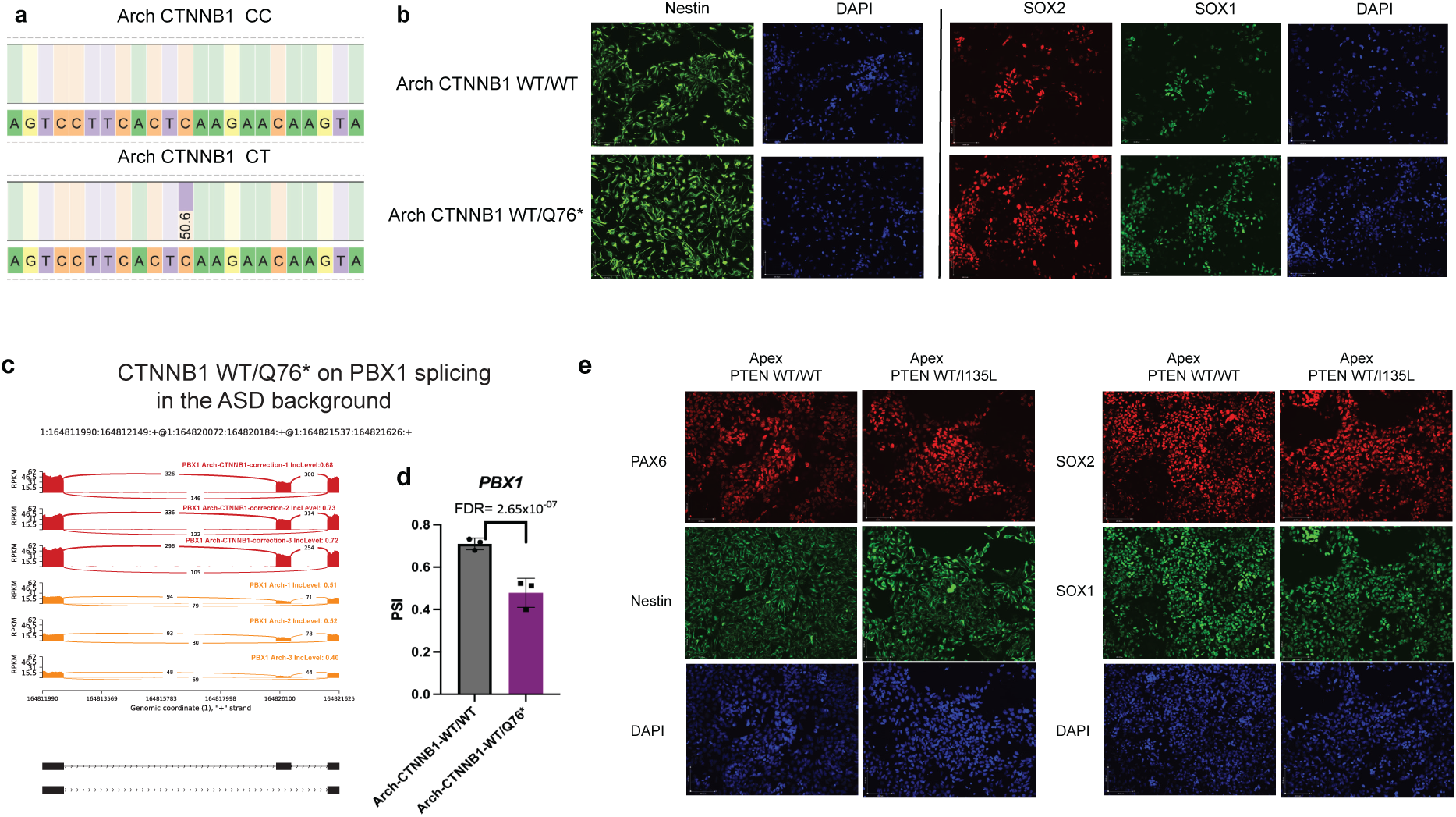
Altered *PBX1* expression in isogenic PTEN and CTNNB1 mutant NPCs in the ASD genetic backgrounds. **a,** Mi-seq confirmation of successful *CTNNB1* mutant correction for ASD Arch iPSC. **b,** Staining for NPC markers in NPCs derived from ASD isogenic *CTNNB1* iPSCs (Arch CTNNB1 WT/WT and Arch CTNNB1 WT/Q76* using the PSC Neural Induction Medium protocol, Scale bar 100µm. **c,** Sashimi Plot visualization for the exon 7 skipping of *PBX1* in Arch CTNNB1 WT/Q76* NPCs. The Y-axis in each plot represents a modified RPKM value. hg38 Chr1:16481990-164821628 spanning PBX1 exon5-8 was shown. Red indicates Arch CTNNB1 WT/WT NPCs, yellow indicates Arch CTNNB1 WT/Q76* NPCs. **d,** rMATS quantification of PSI (percent spliced in) level in (c). N=3 technical replicates for Arch CTNNB1 WT/WT NPCs, N=3 technical replicates for Arch CTNNB1 WT/Q76* NPCs. False discovery rate (FDR) was calculated by rMATS based on the Benjamini-Hochberg approach; error bars represent SD. **e,** Staining for NPC markers in NPCs derived from ASD isogenic PTEN iPSCs (Apex PTEN WT/WT and Apex WT/I135L using the PSC Neural Induction Medium protocol, Scale bar 100µm.

**Extended Data Fig. 4.**
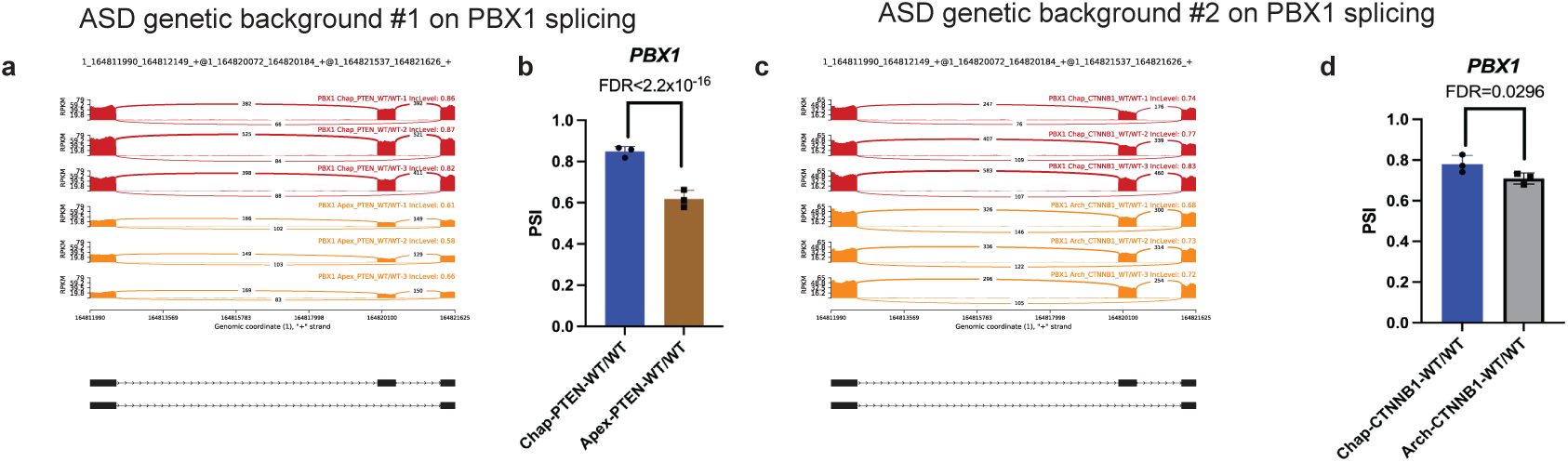
ASD genetic background alters PBX1 alternative splicing in NPCs. **a,** Sashimi Plot visualization for the exon 7 skipping of PBX1 in Apex PTEN WT/WT compared to Chap PTEN WT/WT NPCs. The Y-axis in each plot represents a modified RPKM value. hg38 Chr1:16481990-164821628 spanning PBX1 exon5-8 was shown. Red indicates Chap PTEN WT/WT NPCs, yellow indicates Apex PTEN WT/WT NPCs. **b,** rMATS quantification of PSI (percent spliced in) level in (a). N=3 technical replicates for Chap PTEN WT/WT NPCs, N=3 technical replicates for Apex PTEN WT/WT NPCs. False discovery rate (FDR) was calculated by rMATS based on the Benjamini-Hochberg approach; error bars represent SD. **c,** Sashimi Plot visualization for the exon 7 skipping of PBX1 in Arch CTNNB1 WT/WT compared to Chap CTNNB1 WT/WT NPCs. The Y-axis in each plot represents a modified RPKM value. hg38 Chr1:16481990-164821628 spanning PBX1 exon5-8 was shown. Red indicates Chap CTNNB1 WT/WT NPCs, yellow indicates Arch CTNNB1 WT/WT NPCs. **d,** rMATS quantification of PSI (percent spliced in) level in (c). N=3 technical replicates for Chap CTNNB1 WT/WT NPCs, N=3 technical replicates for Arch CTNNB1 WT/WT NPCs. False discovery rate (FDR) was calculated by rMATS based on the Benjamini-Hochberg approach; error bars represent SD.

**Extended Data Fig. 5.**
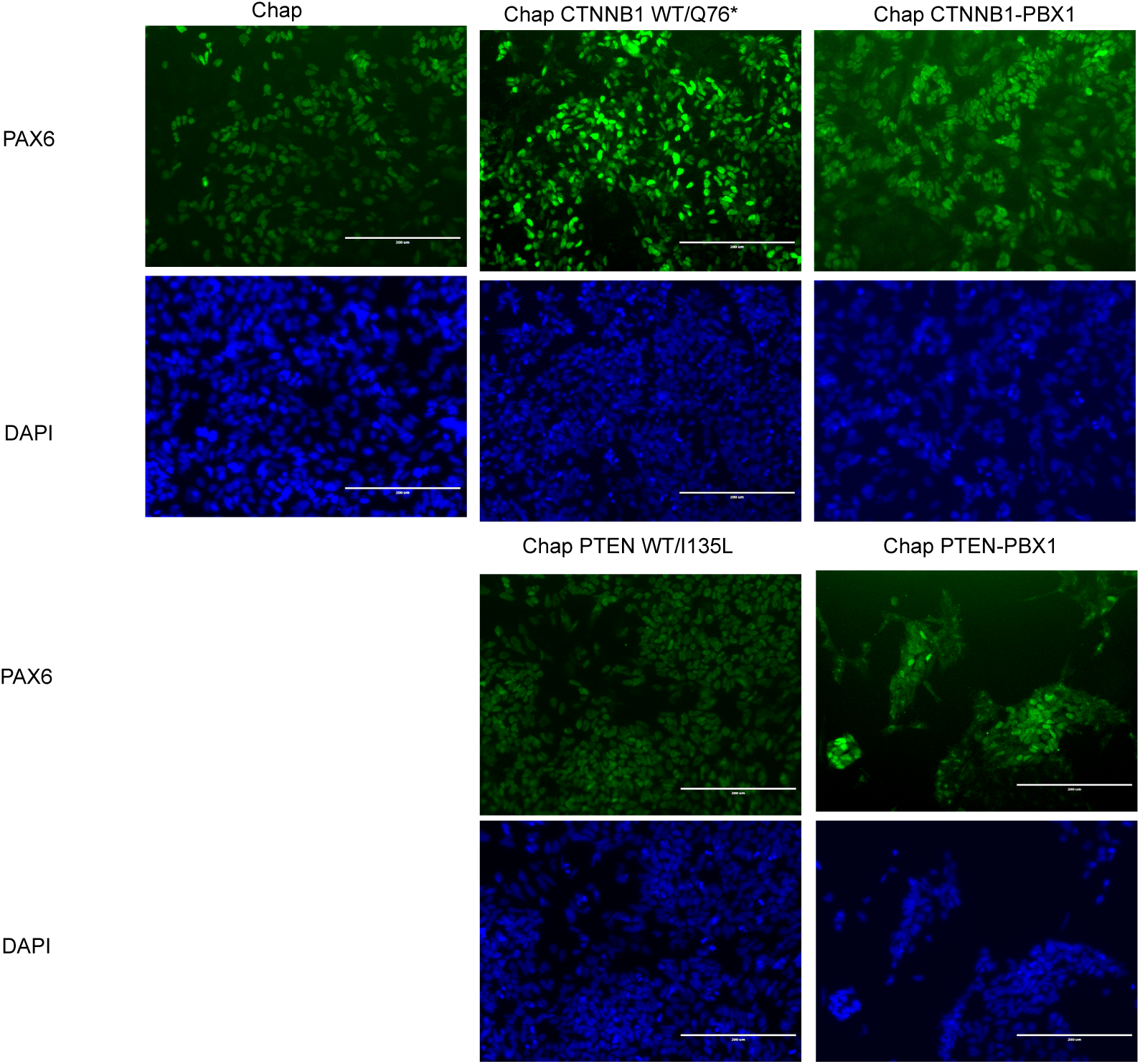
PAX6 staining for isogenic *CTNNB1 PBX1* panel and isogenic *PTEN PBX1* panel iPSC-derived NPCs. Scale bar 200µm.

**Extended Data Fig. 6.**
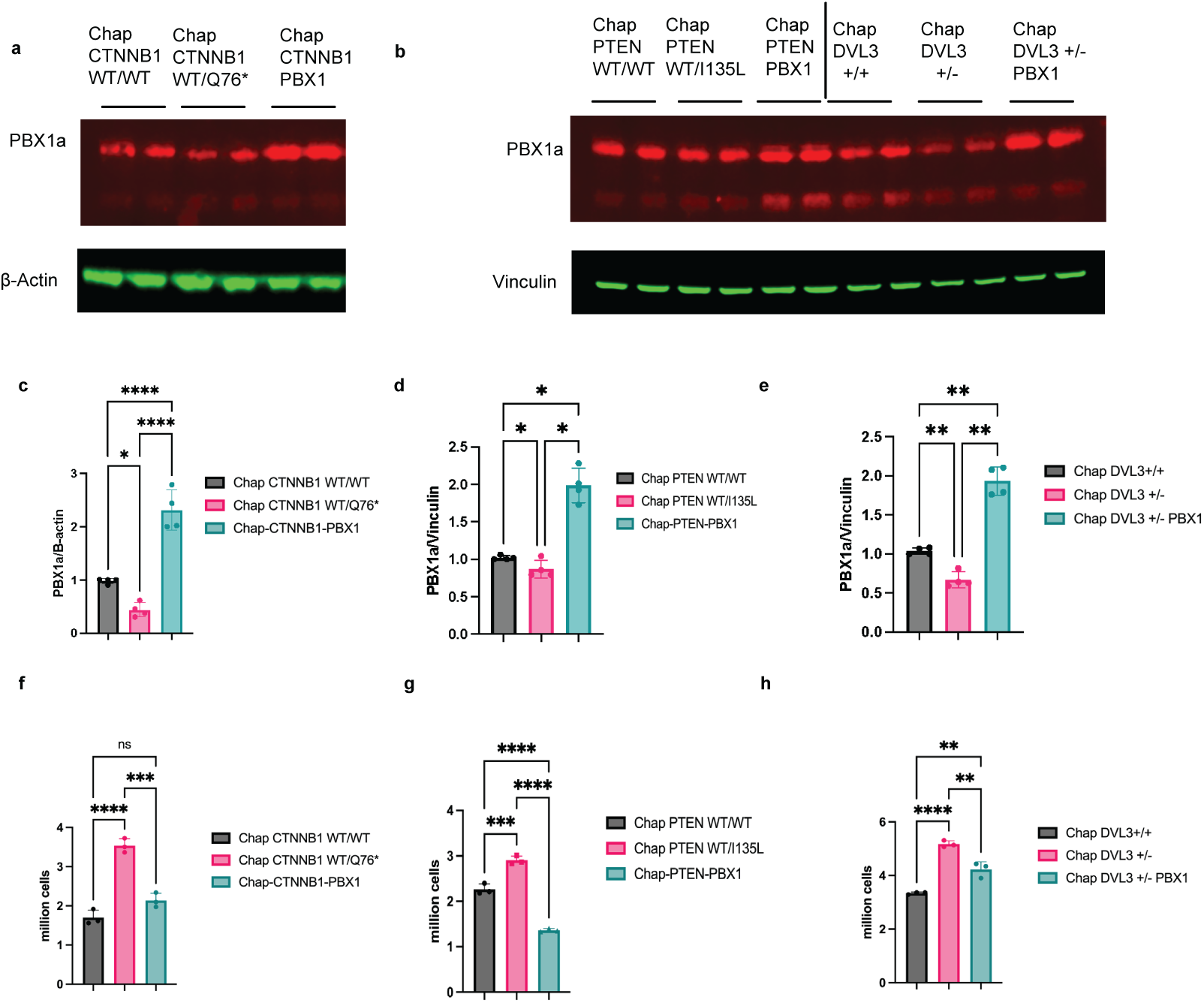
PBX1a overexpression rescues increased NPC proliferation. **a,** Representative western blot probing for PBX1 and β-Actin for protein lysates of isogenic NPC lines chap CTNNB1 WT/WT, Chap CTNNB1 WT/Q76*, and Chap CTNNB1 WT/Q76* overexpressing PBX1a at passage 6. **b,** Representative western blots probing for PBX1 and Vinculin for protein lysates of isogenic NPC lines chap PTEN WT/WT, Chap PTEN WT/I135L, Chap PTEN WT/I135L overexpressing PBX1a at passage 5, chap DVL3^+/+^, Chap DVL3^+/-^, and Chap DVL3^+/-^ overexpressing PBX1a at passage 2. **c,** Quantification of PBX1a / β-Actin in Fig. 4a and Extended Data Fig.6a, p values were calculated with one-way ANOVA with Tukey’s correction for multiple comparisons. *p < 0.05; ****p < 0.0001; error bars represent SD. **d,** Quantification of PBX1a / Vinculin in Fig. 4b and Extended Data Fig.6b, p values were calculated with one-way ANOVA with Holm-Šídák’s correction for multiple comparisons. *p < 0.05; error bars represent SD. **e,** Quantification of PBX1a / Vinculin in Fig. 4c and Extended Data Fig.6b, p values were calculated with one-way ANOVA with Holm-Šídák’s correction for multiple comparisons. **p < 0.01; error bars represent SD. **f,** Cell proliferation assay for isogenic NPC lines Chap CTNNB1 WT/WT, Chap CTNNB1 WT/Q76*, and Chap CTNNB1 WT/Q76* overexpressing PBX1a at passage 6. NPCs were seeded 5x10^5 cells per well of 6-well plate at day 0, and cell counting was performed at day 3. **g,** Cell proliferation assay for isogenic NPC lines Chap PTEN WT/WT, Chap PTEN WT/I135L, and Chap PTEN WT/I135L overexpressing PBX1a at passage 5. NPCs were seeded 5x10^5 cells per well of 6-well plate at day 0, and cell counting was performed at day 3. **h,** Cell proliferation assay for isogenic NPC lines chap DVL3^+/+^, Chap DVL3^+/-^, and Chap DVL3^+/-^ overexpressing PBX1a at passage 2. NPCs were seeded 6x10^5 cells per well of 6-well plate at day 0, and cell counting was performed at day 3.

**Extended Data Fig. 7.**
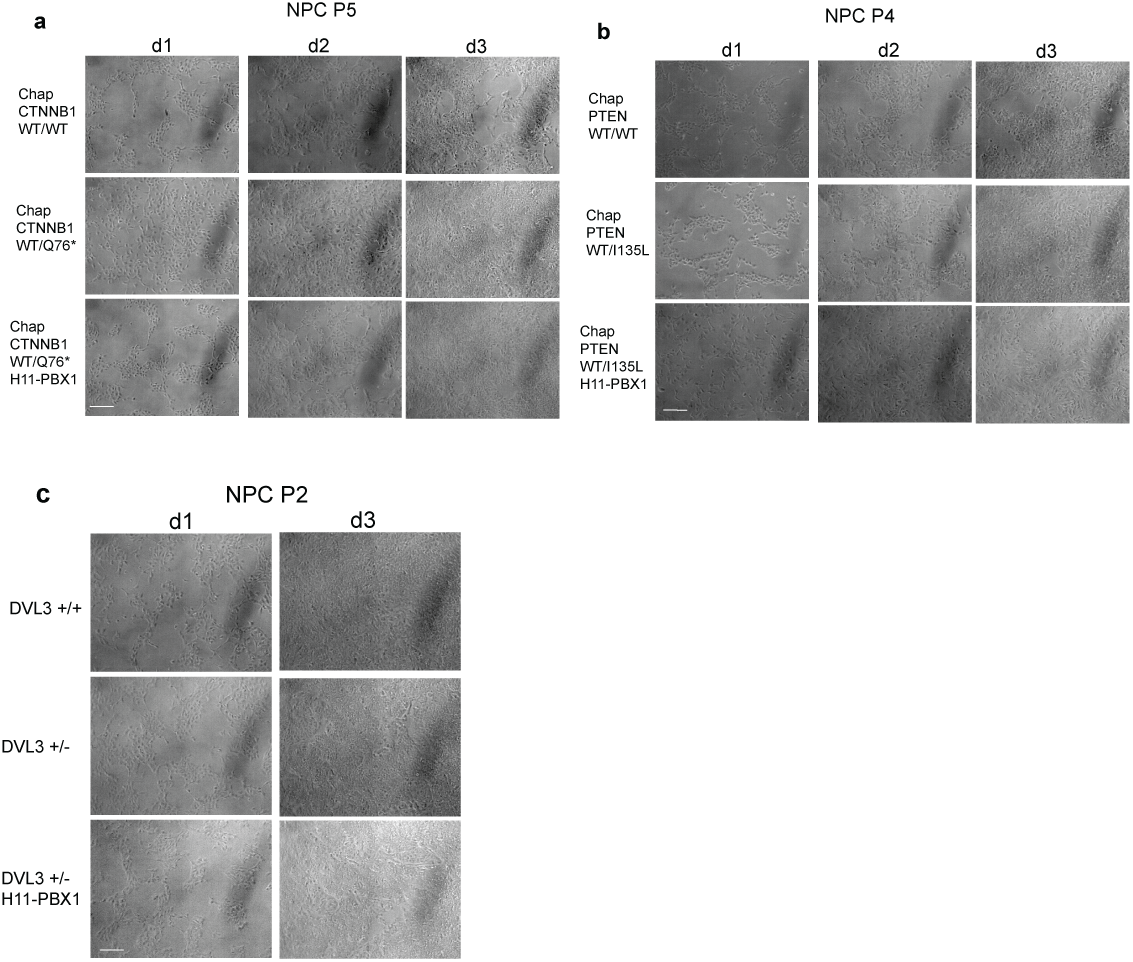
Representative brightfield images for PBX1a rescue experiments. **a,** Representative brightfield images for 2D NPCs Chap CTNNB1 WT/WT, Chap CTNNB1 WT/Q76* and Chap CTNNB1 WT/Q76* overexpressing PBX1a at day1, day2 and day3 of passage 5. Cells were seeded at equal cell numbers for all genotypes. Scale bar: 200 µm. **b,** Representative brightfield images for 2D NPCs Chap PTEN WT/WT, Chap PTEN WT/I135L and Chap PTEN WT/I135L overexpressing PBX1a at day1, day2 and day3 of passage 4. Cells were seeded at equal cell numbers for all genotypes. Scale bar: 200 µm. **c,** Representative brightfield images for 2D NPCs Chap DVL3^+/+^, Chap DVL3^+/-^ and Chap DVL3^+/-^ overexpressing PBX1a at day1 and day3 of passage 2. Cells were seeded at equal cell numbers for all genotypes. Scale bar: 200 µm.

**Extended Data Fig. 8.**
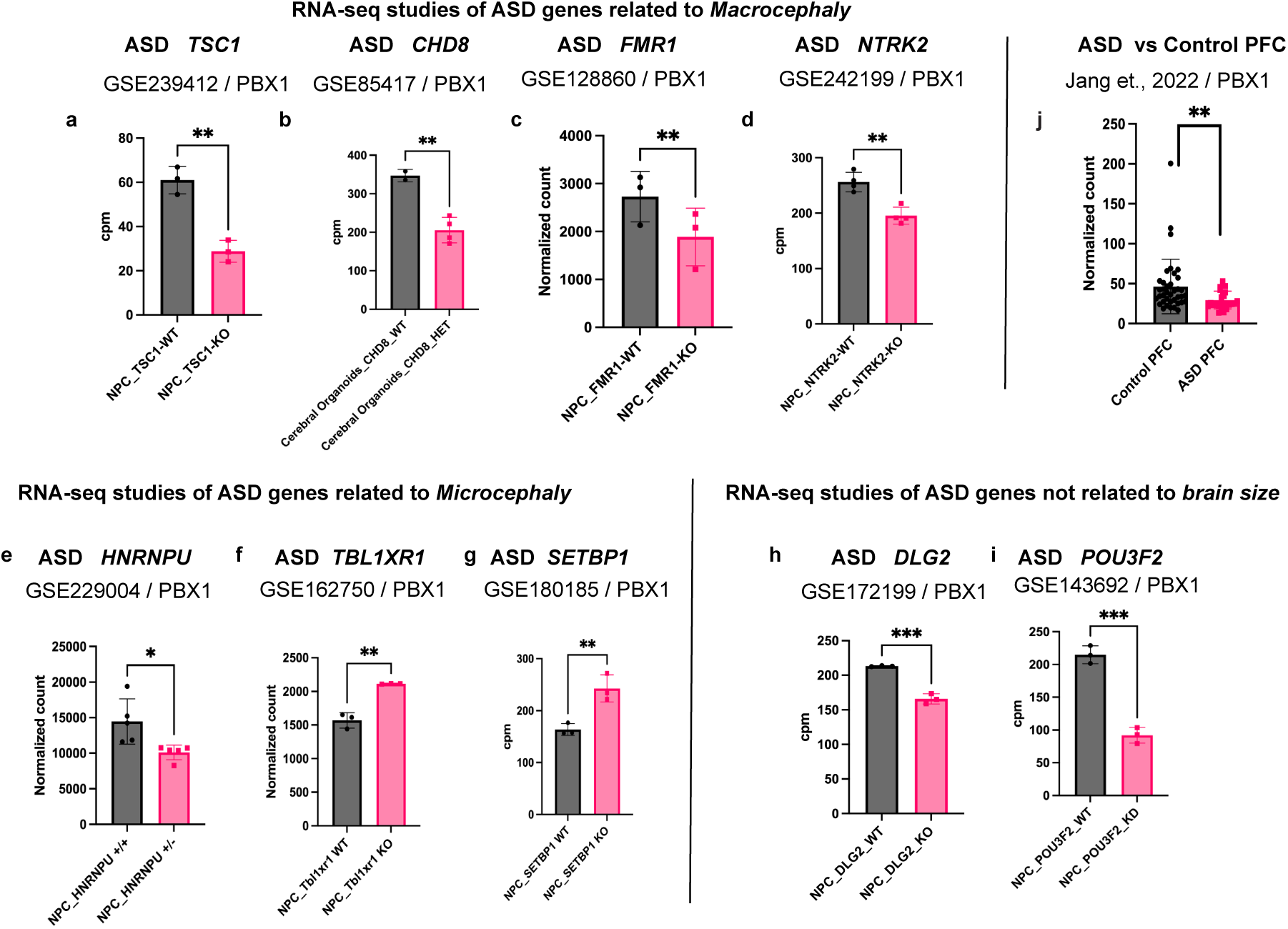
*PBX1* dysregulation identified from other ASD models. RNA-seq studies of ASD genes related to macrocephaly.**a,** RNA-seq quantification for *PBX1* in *TSC1* KO NPCs. N=3 for *TSC1* WT NPCs, N=3 for TSC1 KO NPCs. **p < 0.01; unpaired two-tailed *t* test. **b,** RNA-seq quantification for *PBX1* in CHD8^+/-^ 3D cerebral organoids, cpm means count per million. N=2 for CHD8^+/+^, N=4 for CHD8^+/-^. **p < 0.01; unpaired two-tailed *t* test. **c,** RNA-seq quantification for *PBX1* in FMR1 KO iPSC-derived NPCs. N=3 for *FMR1* WT NPCs, N=3 for *FMR1* KO NPCs. **p < 0.01; unpaired two-tailed *t* test. **d,** RNA-seq quantification for *PBX1* in *NTRK2* KO in the human NPC cell line (ReNcell VM). N=4 for *NTRK2* WT NPCs, N=4 for *NTRK2* KO NPCs. **p < 0.01; unpaired two-tailed *t* test. RNA-seq studies of ASD genes related to microcephaly. **e,** RNA-seq quantification for *PBX1* in *HNRNPU* KO NPCs. N=5 for *HNRNPU*^+/+^ NPCs, N=5 for *HNRNPU*^+/-^ NPCs. *p < 0.05; unpaired two-tailed *t* test. **f,** RNA-seq quantification for *Pbx1* in *Tbl1xr1* KO NPCs. N=3 for *Tbl1xr1* WT NPCs, N=3 for *Tbl1xr1* KO NPCs. **p < 0.01; unpaired two-tailed *t* test. **g,** RNA-seq quantification for *PBX1* in *SETBP1* KO 2D NPCs, cpm means count per million. N=3 for *SETBP1* WT NPCs, N=3 for *SETBP1* KO NPCs. **p < 0.01; unpaired two-tailed *t* test. RNA-seq studies of ASD genes not related to brain size. **h,** RNA-seq quantification for *PBX1* in *DLG2* KO 2D NPCs, cpm means count per million. N=3 for H7 hESC *DLG2* WT NPCs, N=3 for H7 hESC *DLG2* KO NPCs. ***p < 0.001; unpaired two-tailed *t* test. **i,** RNA-seq quantification for *PBX1* in *POU3F2* knockdown (KD) NPCs. N=3 for *POU3F2* WT NPCs, N=3 for *POU3F2* KD NPCs. ***p < 0.001; unpaired two-tailed *t* test. ASD vs Control PFC. **j,** Bulk RNA-seq quantification for *PBX1* in ASD postmortem prefrontal cortex(PFC). N=38 for control PFC, N=25 for ASD PFC. **p < 0.01; unpaired two-tailed *t* test.

